# Parasitoid wasp venoms degrade *Drosophila* imaginal discs for successful parasitism

**DOI:** 10.1101/2024.06.13.598595

**Authors:** Takumi Kamiyama, Yuko Shimada-Niwa, Hitoha Mori, Naoki Tani, Hitomi Takemata-Kawabata, Mitsuki Fujii, Akira Takasu, Minami Katayama, Takayoshi Kuwabara, Kazuki Seike, Noriko Matsuda-Imai, Toshiya Senda, Susumu Katsuma, Akira Nakamura, Ryusuke Niwa

## Abstract

Parasitoid wasps, one of the most diverse and species-rich animal taxa on Earth, produce venoms that manipulate host development and physiology to exploit host resources. However, mechanisms of venom action remain poorly understood. Here, we show that infection of host *Drosophila* by the endoparasitoid wasp, *Asobara japonica*, triggers imaginal disc degradation (IDD) by inducing apoptosis, autophagy, and mitotic arrest, leading to impaired host metamorphosis. A multi-omics approach identified two venom proteins of *A. japonica* necessary for IDD. Knockdown experiments targeting the venom genes revealed that in concert with host immune suppression, IDD is essential for successful parasitism. Our study highlights a venom-mediated hijacking strategy of the parasitoid wasp that allows host larvae to grow, but ultimately kills the hosts.

## Main Text

Parasitoid wasps are one of the most diverse animal groups on Earth, accounting for approximately 20% of insect species (*1*). This taxonomic diversity reflects successful survival parasitism strategies. One group of endoparasitoid wasps injects eggs in hosts to exploit host nutrients. Some parasitoid wasps, termed koinobiont, grow together with their hosts without killing the host immediately (Fig. 1A). To accomplish successful parasitism, koinobiont wasps produce various venom components, including venom proteins, virus-like particles, polydnaviruses, and microRNAs (*2*). These venoms reportedly target host immune systems and/or neuroendocrine systems, protecting wasp eggs and modulating host development to ensure parasitoid development (*3*, *4*). On the other hand, the effect of venoms on other tissues has not been reported. Moreover, venom constituents and their roles in koinobiont parasitism remain poorly understood.

**Fig. 1.**
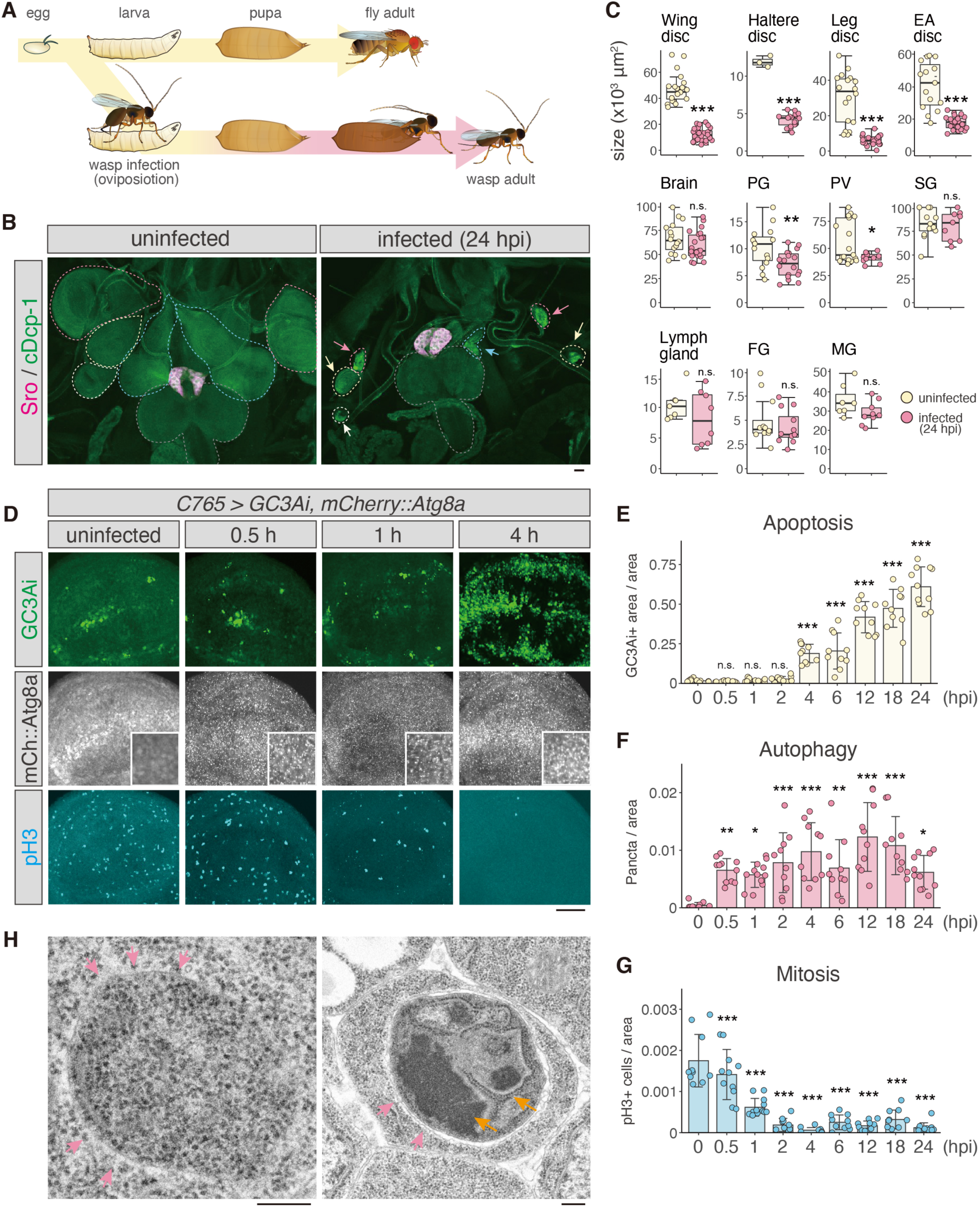
Imaginal disc degradation (IDD) after parasitism by *Asobara japonica*. (**A**) Development of *Asobara japonica*. *A. japonica* deposits an egg inside the fly larval body. The infected fly larva can grow into a pupa, but is completely replaced by a wasp inside the fly pupal case. (**B**) Imaginal disc degradation (IDD) in the infected fly larva. Uninfected and infected (24 hpi) larvae were immunostained for Shroud (Sro, magenta) and cleaved Dcp-1 (cDcp-1, green). Sro labels the prothoracic gland (PG), while cDcp-1 is a marker of apoptosis. Dashed lines and arrows indicate the wing discs (red), eye-antenna discs (blue), haltere discs (white), leg discs (yellow), and brain (grey). Scale bar, 50 µm. hpi, hours post-infection. (**C**) Size of tissues in uninfected (yellow) and infected (red, 24 hpi) larvae. EA, eye-antenna; PV, proventriculus; SG, salivary gland; FG, female gonad, MG, male gonad. (**D**) Time course of apoptosis (GC3Ai, green), autophagy (mCherry::Atg8a, white), and mitosis arrest (pH3, cyan) in uninfected and infected larvae at 0.5, 1, and 4 hpi. Insets of mCherry::Atg8a images show a magnified view of the dorsal-ventral boundary. pH3-positive cells were visualized using anti-pH3 antibody. Scale bar, 50 μm. See also fig. S1E. (**E-G**) Ratio of the GC3Ai-positive area, number of mCherry::Atg8a puncta, number of pH3-positive cells to wing disc area (μm^2^) at each time point (hpi). (**H**) Transmission electron microscopy (TEM) images of autophagosomes and autolysosomes in wing discs at 2 hpi. Double membrane structures are marked with red arrows, while degraded mitochondria and ER membrane are marked with orange arrows. Scale bars, 200 nm. Statistics: (C) Two-tailed Student’s t-test. (E-G) one-way ANOVA followed by Tukey’s multiple comparisons test. *P < 0.05, **P < 0.01, ***P < 0.005; n.s., not significant (P > 0.05). For all bar graphs, means ± standard deviation (SD) of all biological replicates is shown.

To pursue a functional genetic approach to analysing venom components underlying the koinobiont strategy, we focused on the endoparasitoid wasp, *Asobara japonica* Belokobylskij (Hymenoptera: Braconidae) that parasitizes a broad range of *Drosophila* species (Fig. 1A) (*5*, *6*). Among wasps that parasitize the fruit fly, *Drosophila melanogaster*, *A. japonica* has a parthenogenic strain with a high parasitism success rate (*7*, *8*) (fig. S1A). Therefore, we can efficiently monitor developmental processes of koinobionts and host larvae. Moreover, we have previously performed whole-genome sequencing analysis and established a protocol for RNAi methods in *A. japonica*, facilitating the functional analysis of venom genes (*9*).

### Imaginal disc degradation in host flies

To examine how *A. japonica* affects host larval development, we performed infection experiments with third-instar (L3) larvae of *D. melanogaster*. After *A. japonica* oviposited an egg into a fly larva (Movie S1), we dissected host L3 larvae at 24-h post-infection (hpi). Strikingly, adult precursor tissues called imaginal discs were dramatically shrunken in infected L3 larvae, compared to uninfected controls (Fig. 1B, C). In contrast, the central nervous system (the brain and ventral nerve cord) and prothoracic gland (an ecdysteroid-producing organ), which are critical for growth and pupariation (*10*), remained intact in infected larvae. This observation is consistent with the fact that infected host larvae develop into pupae. Comparing tissues between uninfected and infected larvae at 24 hpi, imaginal discs were drastically reduced in size, while other tissues were only slightly reduced, if at all (Fig. 1C). We named this *A. japonica* infection-induced phenotype as imaginal disc degradation (IDD). Moreover, infected L3 larvae delayed pupariation timing compared with uninfected larvae (fig. S1B-D). This delayed phenotype could be attributed to damaged imaginal discs, as reported previously (*11–14*).

### Cellular mechanisms of IDD

To characterize IDD at the cellular level, we focused on wing imaginal discs as representative imaginal discs. We tested the following biological reporters: GC3Ai (an apoptosis reporter), phospho-histone H3 (pH3, a mitosis marker), and mCherry::Atg8a (an autophagy reporter). GC3Ai is a reporter that emits GFP fluorescence upon cleavage by active caspases (*15*). In wing discs of uninfected L3 larvae, only a few cells underwent apoptosis and autophagy, whereas mitotic cells were prominent across tissues (Fig. 1D). In contrast, infected L3 larvae showed substantial induction of apoptosis and autophagy, along with strong suppression of mitosis, within 4 hpi (Fig. 1D-G). These three events were observed at distinct time points post-infection (fig. S1E). Autophagy induction was the earliest event, and distinct punctate signals of mCherry::Atg8a, which reflect autophagosome formation, were detected as early as 0.5 hpi (Fig. 1D, F). Transmission electron microscopy analysis confirmed typical structures of autophagosomes and autolysosomes, which are marked by double membrane-bound vesicles containing undigested cytoplasmic materials and partially degraded organelles (*16*, *17*) (Fig. 1H). They were hardly detected in control wing disc cells from uninfected larvae. Induction of autophagy was followed by mitotic arrest, which was drastically suppressed at 1 hpi (Fig. 1D, G). The GC3Ai signal started to increase at 4 hpi (Fig. 1D, E). At 24 hpi, not only imaginal discs, but also some other tissues, including the lymph gland and female gonads also exhibited apoptosis (fig. S2).

Venom components of *A. japonica* reportedly suppress the host cellular immune defence system (*18*, *19*). Whereas we confirmed that lymph glands were severely damaged at 24 hpi, there was no clear evidence that they were damaged at 4 hpi (fig. S2A-J). This observation suggests that IDD is induced prior to disturbance of the hematopoietic organ. Furthermore, we examined the contribution of hemocytes to IDD by genetically ablating hematopoietic organs and hemocytes in host larvae with forced expression of proapoptotic genes. IDD persisted in hemocyte-free hosts, supporting the hypothesis that host hemocytes were not involved (fig. S3). These results also suggest that *A. japonica* infection-induced IDD is not mediated by the immune system, but rather by acute cell death and mitotic arrest in imaginal discs.

*A. japonica*-induced apoptosis was caspase-dependent, as cleaved *Drosophila* Death caspase-1 (cDcp-1) signals were suppressed in host larvae expressing *p35*, which encodes a baculovirus-derived caspase inhibitor (*20*). Autophagy and mitotic arrest were not suppressed in *p35*-expressing host larvae (fig. S4A-F), suggesting that autophagy and mitotic arrest are regulated independently of the apoptotic signalling pathway. Upon simultaneously overexpressing *p35* and *Rubicon*, which encodes a negative regulator of autophagy (*21*), both apoptosis and autophagy were inhibited, but mitotic arrest persisted in wing discs of infected host larvae (fig. S4G-J). Consistent with these observations, wing disc size was not restored in *p35* and *Rubicon* double-expressing flies (fig. S4K, L). Taken together, our results suggest that *A. japonica* infection-induced apoptosis, autophagy, and mitotic arrest are regulated independently.

### IDD is linked to host death

To further explore the relationship between IDD and parasitism, we examined imaginal discs of the non-host *Drosophila* species, *D. elegans* and *D. ficusphila* (*7*). *A. japonica* injected eggs and venom into these species; however, its offspring failed to emerge from host pupal cases (Fig. 2A, F). In *D. elegans,* a host fly did not eclose from the pupal case, resulting in death of both host and parasitoid (Fig. 2A). Interestingly, apoptosis, autophagy, and mitotic arrest were detected in wing discs, suggesting that IDD was induced in *D. elegans* (Fig. 2B-E). Effects of infection on *D. elegans* will be further examined and discussed later.

**Figure 2:**
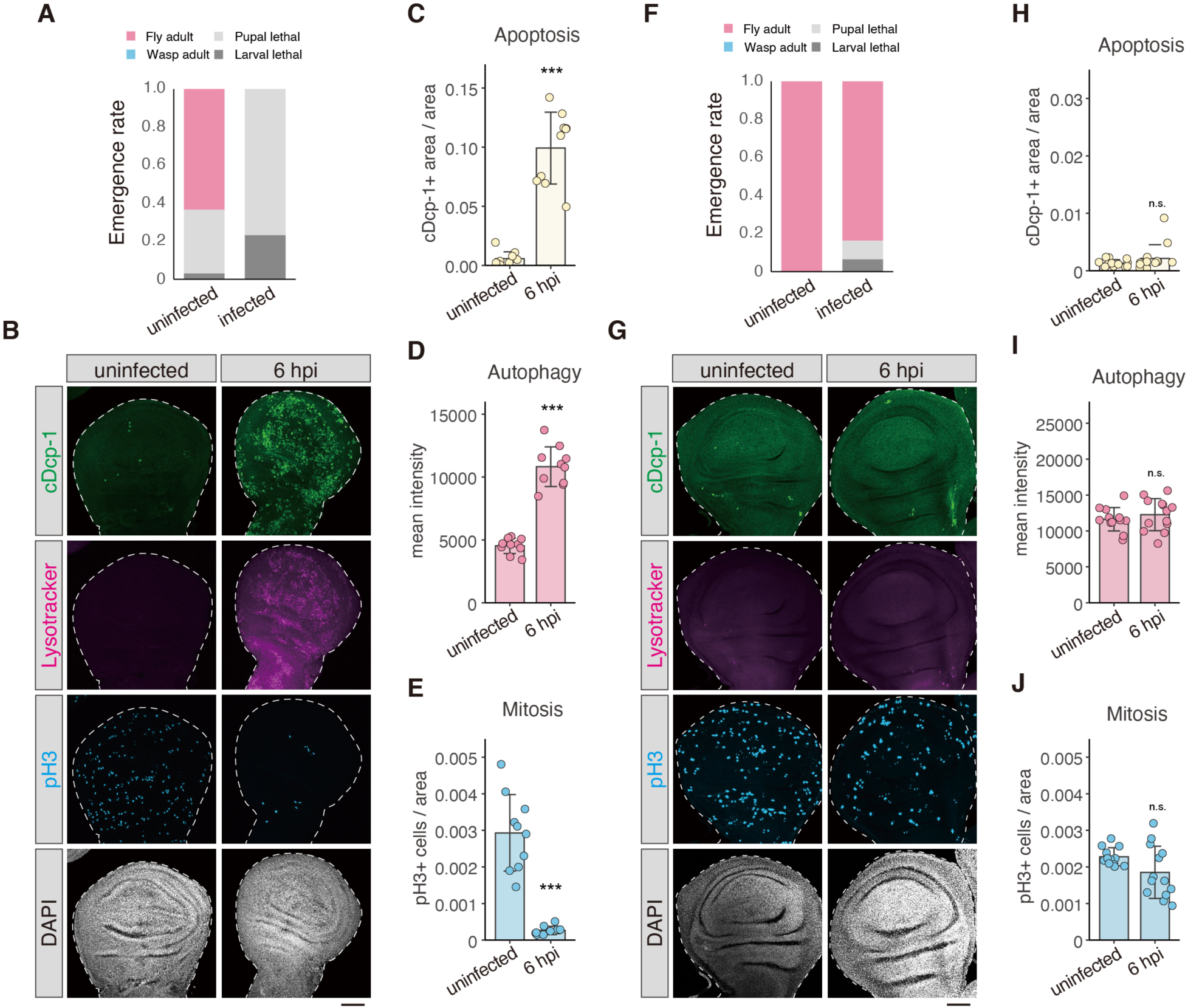
IDD is linked to host death in *D. elegans*, but not in *D. ficusphila*. (**A, F**) Parasitism success rates of *D. elegans* and *D. ficusphila* (each n = 30). (**B, G**) Wing discs from uninfected and 6 hpi fly larvae. Apoptosis and mitosis were detected using anti-cDcp1 (green) and anti-pH3 (blue) antibodies, respectively. Autophagy was visualized with LysoTracker (magenta). Nuclei were stained with DAPI (white). Scale bar, 50 µm. (**C-E, H-J)** Ratio of cDcp-1-positive area (C, H), mean intensity of LysoTracker signal (D, I), number of pH3-positive cells (E, J) to wing disc area (μm^2^) from uninfected and 6 hpi fly larvae. For all bar graphs, means ± standard deviation (SD) for all biological replicates is shown. Statistics: Two-tailed Welch’s t-test. *P < 0.05, **P < 0.01, ***P < 0.005; n.s., not significant (P > 0.05).

Unlike *D. elegans*, infected *D. ficusphila* larvae grew normally to adulthood (Fig. 2F), indicating complete resistance to attack by *A. japonica*. IDD was not induced in imaginal discs of *D. ficusphila* (Fig. 2G-J), suggesting that IDD is associated with host death. We speculate that IDD is the key to high parasitism success in *A. japonica*, preventing host metamorphosis after pupariation.

### Venom gland components induce IDD

We reasoned that IDD was induced by venom components and/or an egg of *A. japonica*, both of which are injected into the host hemocoel during oviposition (*18*, *22*). To examine whether venom and/or eggs have IDD-inducing activity, we performed microinjection experiments, as described previously (*18*). We dissected *A. japonica* adult females to collect venom glands and ovaries separately and prepared lysates of each (Fig. 3A-C). Microinjection of phosphate-buffered saline (PBS control) or ovary lysate had no effect on wing discs (Fig. 3D-G). In contrast, microinjection of the whole-body or venom gland lysate led to apoptosis, autophagy, and mitotic arrest in wing discs of naïve test larvae. To further examine whether venom gland-derived substance(s) could act on wing discs directly, we performed an *ex vivo* co-culture assay in which venom glands and wing discs were co-cultured in culture media for 2 h. Compared with controls, wing discs cultured with venom glands exhibited increased apoptosis, autophagy, and mitotic arrest (fig. S5). Notably, IDD-inducing activity was detected even when venom glands were not crushed. These results suggest that venom components secreted from venom glands of *A. japonica,* act directly on *Drosophila* wing discs.

**Fig. 3:**
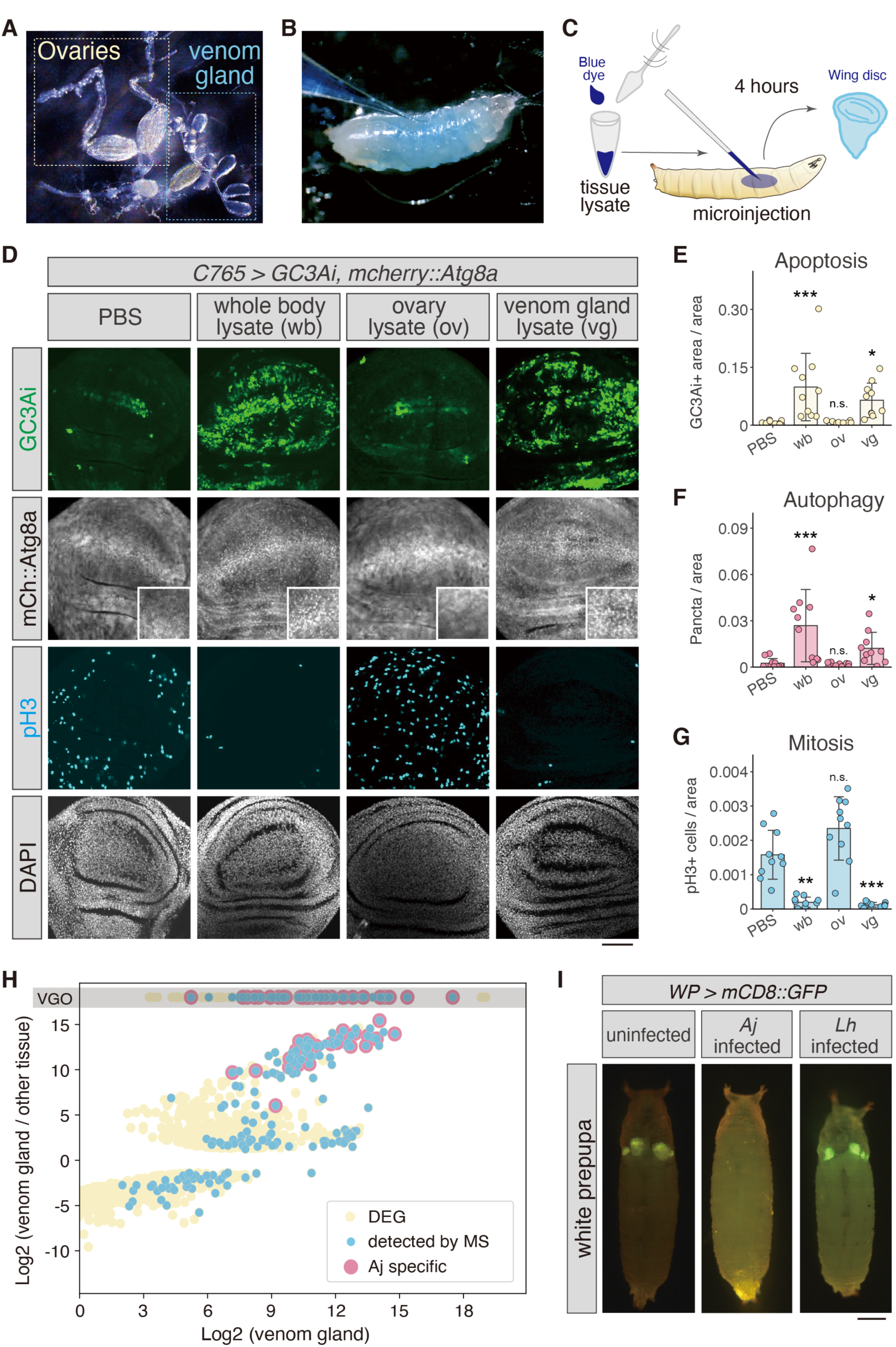
IDD-inducing activity comes from the venom gland of *A. japonica*. (**A**) *A. japonica* ovaries and venom gland. (**B**) Microinjection of tissue lysate into uninfected fly larvae. (**C**) Scheme of the microinjection experiment. Wasp tissue lysate was injected to uninfected fly larvae. Wing discs were dissected 4 h after injection. (**D**) GC3Ai (green), mCherry::Atg8a (white), pH3 (cyan), and DAPI (white) in wing discs of fly larvae 4 h after injection of phosphate-buffered saline (PBS), whole body lysate (wb), ovary lysate (ov), or venom gland lysate (vg). Insets of mCherry::Atg8a images are magnified views of the dorsal-ventral boundary. Scale bar, 50 µm. (**E-G**) Ratio of GC3Ai positive area, number of mCherry::Atg8a puncta, number of pH3-positive cells to wing disc area (μm^2^) of fly larvae in each condition. n=10 in each sample. (**H**) Scatterplot of differentially expressed genes (DEGs, yellow dots) from RNA-sequencing analysis. The X-axis represents log2-scaled values of expression in the venom gland, while the Y-axis is relative expression values in the venom gland to the other tissues, respectively. VGO (venom gland only) indicates zero read count in the other tissue sample. Proteome-detected genes and *A. japonica-*specific genes via comparative genomics are labelled with blue dots and red dots, respectively. See also Data S1. (I) Uninfected, *A. japonica*-infected, and *L. heterotoma*-infected flies at the prepupal stage. Wing and haltere discs are visualized with GFP fluorescence (genotype: *WP>mCD8::GFP*). Scale bar, 1 mm. For all bar graphs, mean ± standard deviation (SD) for all biological replicates is shown. Statistics: one-way ANOVA followed by Bonferroni’s multiple comparisons test. *P < 0.05, **P < 0.01, ***P < 0.005; n.s., not significant (P > 0.05).

### Identification of IDD factors

To identify *A. japonica* venom components crucial for IDD-inducing activity, we conducted RNA-sequencing (RNA-seq) analysis based on our gene annotation database (*9*). We compared expression levels of 12,508 annotated genes between the venom gland and other parts of the wasp body, which led to identification of 807 genes that were significantly enriched in venom glands (Fig. 3H, Data S1). Subsequently, we conducted a proteome analysis of venom gland lysate and detected 197 gene products among 807 genes in the venom gland. To further focus on IDD-related genes among the 197 candidates, we conducted comparative genomics against another parasitoid, *Leptopilina heterotoma*. Although this species parasitizes *Drosophila* larvae and a single adult wasp ecloses from the host pupal case, as in *A. japonica*, IDD was not induced after infection (Fig. 3I). Thus, we speculated that venom components crucial for IDD-inducing activity are not conserved in the *L. heterotoma* genome. Based on this assumption, we focused on 63 “*Aj*-specific” genes, which were determined using tblastn E-value > 0.001 as a threshold (Fig. 3H, Data S1). More than 70% (52 genes) of *Aj*-specific genes encoded predicted secretory proteins with signal peptide sequences in their N-terminal regions. On the other hand, there were no known enzymes involved in specific metabolic or redox pathways.

We then conducted functional screening of IDD-related *A. japonica* genes using a previously established RNAi technique (*9*). In the first screen, 63 genes were divided into two to four gene groups according to their high false discovery rates in the transcriptome analysis. Subsequently, dsRNA mixtures corresponding to the two to four target genes were microinjected into wasp bodies to knock down multiple genes simultaneously. After eclosion, RNAi adult wasps were allowed to parasitize fly L3 larvae expressing the *GC3Ai* reporter in their wing pouch regions (*WP>GC3Ai*). Then we measured the fluorescence intensity of GC3Ai in host larvae, quantifying the apoptosis-inducing activity of RNAi wasps. Following infection, control (*GFP-*RNAi) wasps increased GC3Ai fluorescence in host wing discs at 6 hpi (fig. S6A). In contrast, Group #3 and #5 RNAi most dramatically suppressed the increase of GC3Ai signals.

In the second screening, RNAi wasps were generated to target individual genes in Groups #3 and #5 (Fig. 4A). We eventually identified *gene007424* (Group #3) and *gene008061* (Group #5), both of which encode putative secretory proteins (Fig. 4B). Interestingly, both proteins contain the same domain structure, classified as Domain of Unknown Function 4803 (DUF4803) (*23*, *24*) and seem to exhibit similar three-dimensional structures, as predicted with AlphaFold 2 (Fig. 4B; fig. S6B). RNA-seq analysis using RNA of venom glands confirmed that expression levels of these target genes were significantly reduced in RNAi wasps (Fold changes: 0.067 for *gene007424*, 0.058 for *gene008061*; fig. S6C, D; Data S2). Following knockdown of *gene007424* or *gene008061*, the GC3Ai signal was barely detected in host wing discs. Moreover, autophagy was significantly decreased by either of these RNAi wasps, whereas mitotic arrest was recovered only by *gene007424* RNAi wasps (Fig. 4C-F). We confirmed the similar phenotype with another target region of RNAi, supporting that the phenotype was not due to the off-target effects (fig. S7). Importantly, a substantial restoration of wing disc size resulted from each venom gene knockdown (Fig. 4G, H). Simultaneous knockdown of *gene007424* and *gene008061* did not alter wing disc size when compared with single-gene RNAi, suggesting that these two proteins do not have an additive effect, possibly acting on the same pathway (Fig. 4G, H). We named *gene007424* and *gene008061* as *Imaginal disc degradation factor 1* (*IDDF-1*), and *IDDF-2*, respectively.

**Fig. 4:**
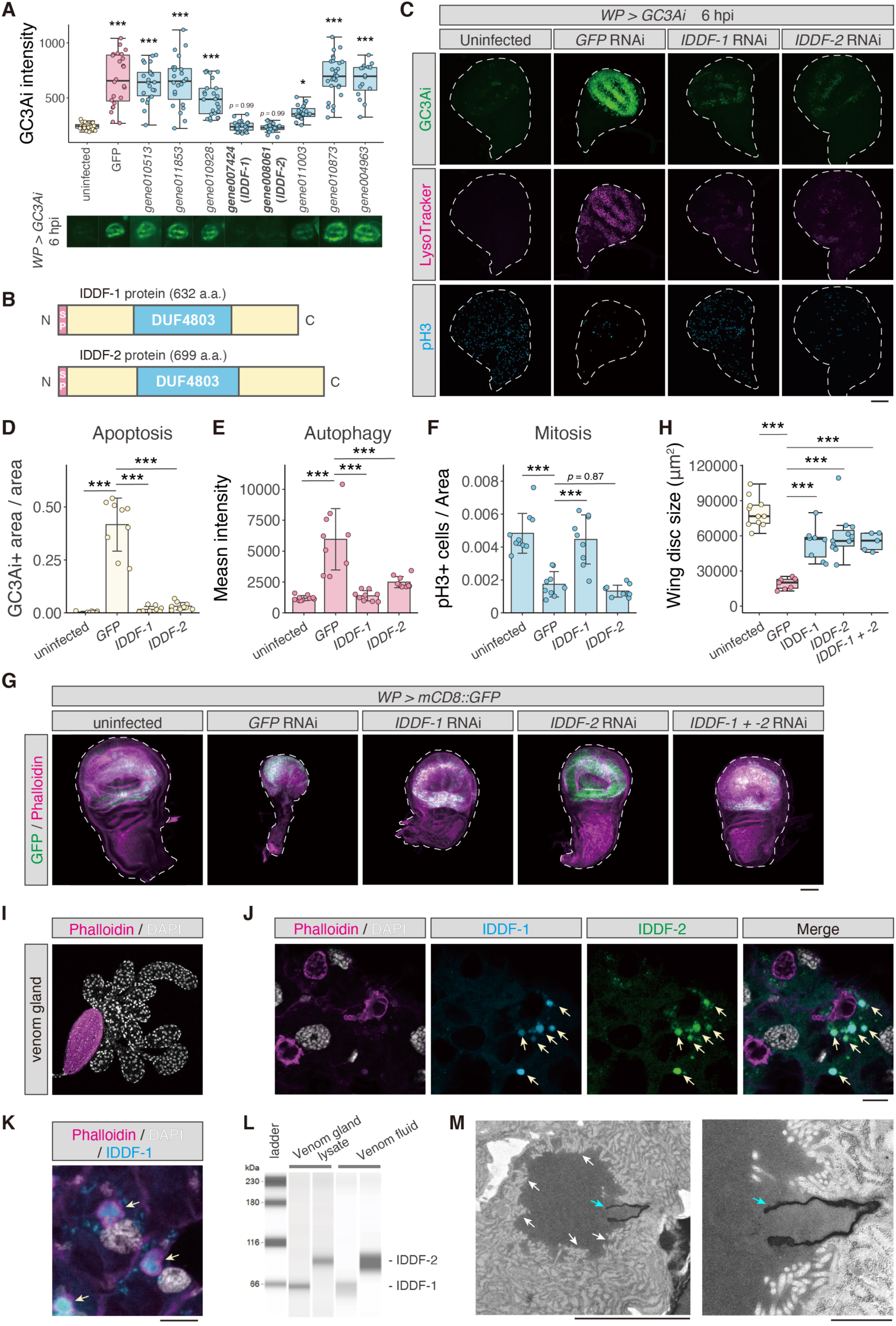
Identification of wasp venom proteins required for IDD. (**A**) RNAi screening against candidate *A. japonica*-specific genes. Fly larvae expressing *GC3Ai* in the wing pouch region (*WP>GC3Ai*) were used as hosts for RNAi wasps. Fluorescence of GC3Ai increased by infection of control (*GFP*-RNAi) wasps compared to that of uninfected larvae. Box-and-whisker plots representing all biological replicates of GC3Ai intensity. In the Group #3 and #5, knockdown of *gene007424* (*IDDF-1*) or *gene008061* (*IDDF-2*) decreased the GC3Ai signal in wing discs. See also fig. S6A. (**B**) Domain structures of IDDF-1 and IDDF-2 proteins. SP, signal peptide sequence; a.a., amino acids; DUF, domain of unknown function. (**C**) Representative images of GC3Ai (green), LysoTracker (magenta), and pH3 (cyan) in wing discs of fly larvae at 6 hpi, infected by *GFP-*, *IDDF-1-*, and *IDDF-2-*RNAi wasps. Scale bar, 50 µm. (**D-F**) Ratio of cDcp-1 positive area, mean intensity of LysoTracker signal, number of pH3-positive cells to wing area (μm^2^) from fly larvae at 6 hpi, infected by *GFP-*, *IDDF-1-* or *IDDF-2-*RNAi wasps. (**G, H**) Restoration of wing disc size by *IDDF-1-* and/or *IDDF-2*-RNAi. Representative images (G) and size (H) of wing discs from uninfected or infected flies at the prepupal stage. Host flies were infected by *GFP-*, *IDDF-1-* and/or *IDDF-2-*RNAi wasps. Wing discs were labelled with GFP (green) and stained with phalloidin (magenta). Scale bar, 50 µm. (**I**) The *A. japonica* venom gland was stained with phalloidin (magenta) and DAPI (white). A rugby ball-like structure is rich in muscles, connected to an ovipositor. (**J, K**) Subcellular co-localization of IDDF-1 and IDDF-2 in the venom gland (yellow arrows). The venom gland was immunostained with anti-IDDF-1 antibody (cyan) and anti-IDDF-2 antibody (green) as well as phalloidin (magenta) and DAPI (white) staining. (**L**) Western blotting analysis of IDDF proteins detected in both venom gland lysate and venom fluid samples. (**M**) Ultrastructure of the venom gland. White arrows indicate microvilli in a circle connecting to a duct-like cuticle (blue arrow). Scale bar, 5 µm, 1 µm. Statistics: one-way ANOVA followed by Dunnett’s test (A). one-way ANOVA followed by Tukey’s multiple comparisons test (D-F, H). *P < 0.05, **P < 0.01, ***P < 0.005; n.s., not significant (P > 0.05).

### Localization of IDDFs in the venom gland

Antibodies were generated to visualize the IDDF proteins using immunostaining and western blotting (Fig. 4I-L). Particle-like signals for IDDF-1 and IDDF-2 were detected in venom gland cells, surrounding actin-rich ring structures (Fig. 4J). Interestingly, particle size tended to increase with wasp age after eclosion (fig. S8A). These signals were not observed in *IDDF-1* or *IDDF-2-*RNAi wasps, so that they reflect endogenous venom proteins (fig. S8B, C). These particles accumulate surrounding or inside the actin-rich ring structures of three-day-old or older wasps (Fig. 4J, K; fig. S8A). Similar actin-rich structures have been described in certain wasp venom glands termed long glands (*25*). In ultrastructural analysis, actin-rich microvilli of venom gland cells surrounded duct-like cuticles, representing presumptive exit sites of venom secretion (Fig. 4M). Importantly, IDDF proteins were detected in venom fluid leaking from tips of wasp ovipositors (Fig. 4L; fig. S8D). Therefore, these venom proteins are probably secreted into the lumen of the gland and injected into the host hemocoel.

To test whether IDDF proteins were sufficient to induce IDD, we generated recombinant proteins with an *in vitro* translation system and performed microinjection experiments for bioassay. Recombinant proteins were successfully detected with antibodies by western blotting (fig. S9A, B). When either IDDF-1 or IDDF-2 protein was injected into naive fly larvae, apoptosis, autophagy, and mitotic arrest were hardly detected, similar to PBS injection and control bacterial protein injection (fig. S9C-F). In contrast, microinjection of both IDDF proteins led to a mild increase in apoptosis and autophagy. This result implies that IDDF-1 and IDDF-2 act cooperatively on host wing discs. On the other hand, neither protein affected mitosis. Moreover, the effect of these recombinant proteins on apoptosis and autophagy was not as prominent as that of whole wasp lysate. This result may reflect lower IDD-inducing activities of recombinant proteins compared to those of endogenous IDDF proteins. Another intriguing possibility is that there remain other venom factors in the venom gland.

### IDDFs ensure success of parasitism

To evaluate whether these IDDFs contribute to parasitism success of *A. japonica,* we examined the parasitism success rate of control and RNAi wasps. In *GFP*-RNAi wasps, more than 80% of pupae led to wasp eclosion (Fig. 5A), which was comparable with that of wild-type wasps (fig. S1A). The remaining 20% proved lethal, preventing emergence of adult flies. In contrast, *IDDF-1*- or *IDDF-2*-RNAi wasps allowed eclosion of a small, but significant number of adult host flies (Fig. 5A), suggesting that *IDDF-*mediated IDD contributes to successful parasitism. However, it should be noted that the parasitism success rate was still high (∼80 %), implying that some other factor is required for successful parasitism of *A. japonica*.

**Fig. 5:**
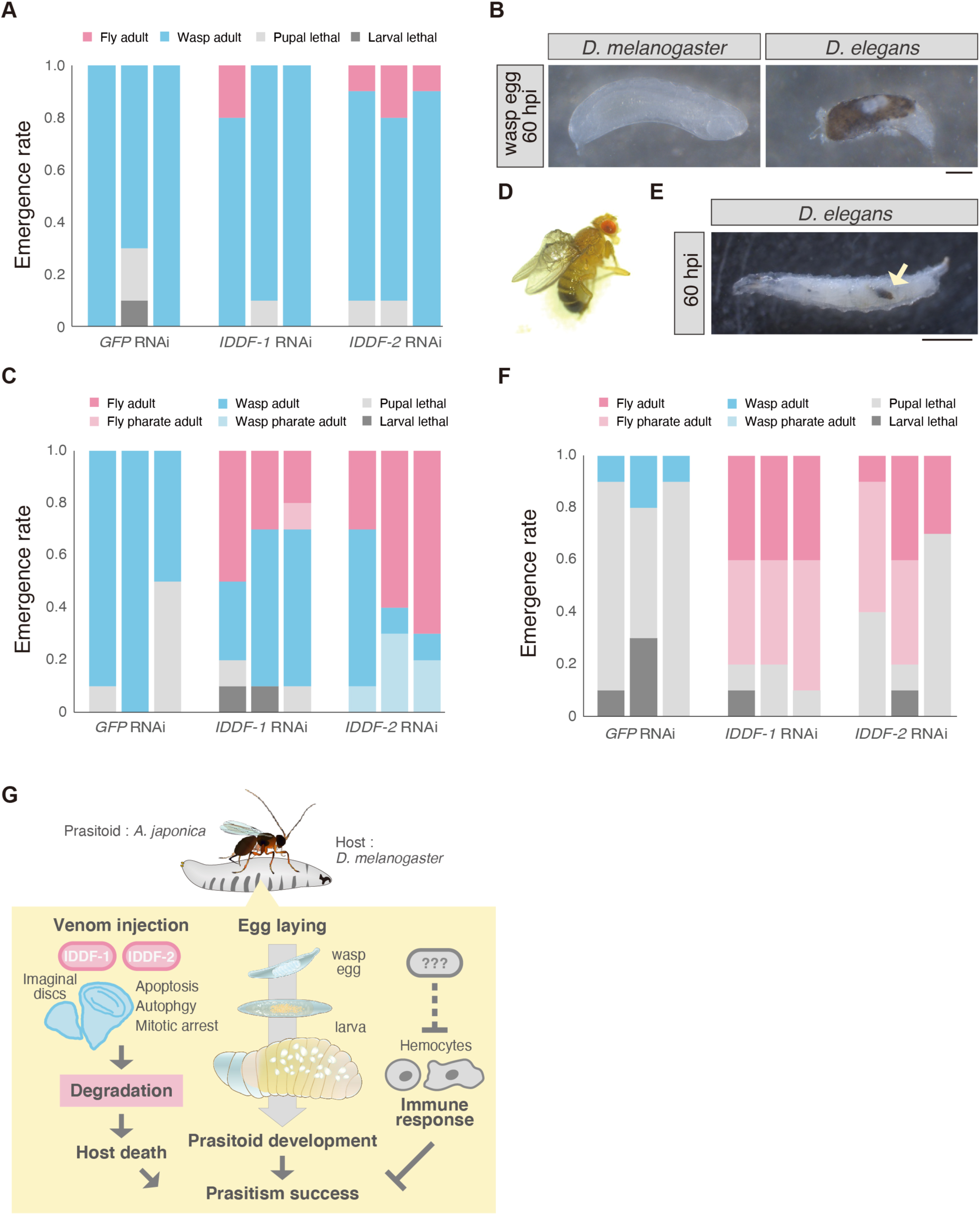
Both IDD and immunosuppression are required for successful parasitism. (**A, C, F**) Parasitism success rate of *A. japonica* on host *D. melanogaster* wild type (*Oregon R*) (A), *tuSz^1^*strain (C), and *D. elegans* (F). A single bar represents the result of a single RNAi wasp, targeting *GFP*, *IDDF-1* or *IDDF-2.* A wasp lays an egg in each of 10 host flies. (**B**) *A. japonica* egg taken from *D. melanogaster* and *D. elegans* host larvae at 60 hpi. Scale bar, 0.1 mm. (**D**) A “survivor” *tuSz^1^* fly infected by an *IDDF-1-*RNAi wasp, whose wings showed a morphological defect. (**E)** Melanized *A. japonica* body (yellow arrow) in the *D. elegans* larva at 60 hpi. Scale bar, 1 mm. (**G**) Schematic of a working model. IDDFs are essential for induction of apoptosis, autophagy, and mitotic arrest-mediated imaginal disc degradation (IDD), ensuring successful parasitism of *A. japonica*. In addition, some unknown venom factors suppress host immune responses that would otherwise prevent successful parasitism.

### IDD and immune suppression

We assumed that the high parasitism success of *A. japonica* requires not only IDD, but also suppression of the fly immune system. Generally, to eliminate wasp eggs, *Drosophila* immune cells (lamellocytes) encapsulate and melanize the eggs (*26*, *27*). However, when wild-type *D. melanogaster* was infected with *A. japonica,* encapsulation of wasp eggs was not observed (Fig. 5B), suggesting that the immune response was actively suppressed. Thus, we speculate that the relatively high parasitism success of *IDDF-*RNAi *A. japonica* may be because suppression of the immune system remained unchanged. To test this hypothesis, we used two types of hosts with high immune activities.

One of these was the *D. melanogaster tumour* (*1*)*Suzuki* (*tuSz^1^*) mutant strain. *tuSz^1^* larvae exhibit moderate constitutive activation of the innate immune system without parasitoid wasp infection owing to the presence of a gain-of-function mutation in the *hopscotch* gene (hop), the *Drosophila* Janus Kinase (JAK) orthologue (*28*). When *GFP*-RNAi wasps were infected with *tuSz^1^* larvae, adult host flies never eclosed (Fig. 5C). In contrast, the eclosion rate of adult host flies increased by up to 50% after infection by *IDDF-1-* or *IDDF-2*-RNAi wasps. Notably, these “survivor” flies exhibited some morphological defects on their wings and legs, possibly due to partial IDD (Fig. 5D).

The other host was wild-type *D. elegans* larvae, which could melanize *A. japonica* eggs upon infection, as mentioned earlier, indicating a more active immune response than that of *D. melanogaster* (Fig. 5B, E). Upon infection by *GFP-*RNAi wasps, similar to wild-type wasps, no *D. elegans* reached adulthood. In contrast, more than 50% of *D. elegans* larvae reached the pharate adult stage or adulthood after *IDDF-1 or IDDF-2-*RNAi wasp infection (Fig. 5F). Collectively, these results strongly support the hypothesis that with a combination of host immune suppression, *IDDF-*mediated IDD is essential for *A. japonica* parasitism with a high success rate (Fig. 5G). This hypothesis is also supported by the fact that *D. ficusphila,* which is completely resistant to *A. japonica* infection, encapsulates wasp eggs and does not exhibit IDD (Fig. 2F-J; fig. S10A).

## Discussion and future perspectives

Our study highlights a novel strategy for the koinobiont endoparasitoid wasp, *Asobara japonica*, in which secretory venom proteins, IDDF-1 and IDDF-2, mediate IDD without disturbing the host neuroendocrine system. We propose that these venom actions lead to apoptosis, autophagy, and mitotic arrest in imaginal discs, thereby blocking host imaginal development. IDD is coupled with host death after metamorphosis.

Mechanisms by which IDDF proteins exert their action on imaginal discs remain unknown. Several lines of evidence have revealed that wasp-induced apoptosis is mediated by the proapoptotic gene *reaper (rpr),* accompanied by a typical cellular response including DNA damage and reactive oxygen species (*29*) (fig. S11A-E). Expression of *rpr* is reportedly upregulated by X-ray irradiation (*30*), indicating that infection by *A. japonica* evokes a strong stress response in imaginal discs. Conversely, wasp-induced apoptosis was distinct from the response to X-ray irradiation, given its induction in polyploid cells (fig. S11F). In future investigations, we plan to investigate how wasp venom components are received by imaginal disc cells, leading to cooperative induction of apoptosis, autophagy, and mitotic arrest. Although a previous study assumes that parasitoid virulence factors are virus-like (*22*), our ultrastructural analysis did not find virus-like structures in venom glands of *A. japonica* (Fig. 4M).

Our data suggest that IDDF-mediated IDD ensures successful parasitism in combination with infection-induced immunosuppression in *D. melanogaster.* When *A. japonica* infects *D. elegans* larvae, IDD is induced, but the host immune system is active on *A. japonica* egg, resulting in the death of both host flies and parasitoid wasps (Fig. 2A, 5B, E; fig. S10A). In both species, RNAi-mediated suppression of IDD increased the adult fly survival rate, suggesting that the IDD-induction mechanism is distinct from suppression of the host immune system. Immune suppression is possibly regulated by some of the other approximately 200 venom proteins in venom glands of *A. japonica*. A recent study using *L. heterotoma* identified *Lar* as an essential regulator of the host immune system. Interestingly, we found that this *Lar* gene is not conserved in *A. japonica*. Therefore, in future studies, it will be interesting to identify *A. japonica* venom proteins that inhibit host immunity. Considering that *D. ficusphila* is resistant to these venoms, it will be also intriguing to compare these three species to determine at the molecular level how susceptibility and resistance to venom proteins have been acquired and modified during the evolutionary “arms race” of host and non-host *Drosophila* species. These studies will also shed a light on *A. japonica* as a potential pest control reagent against *D. suzukii,* a serious economic threat to soft summer fruit, particularly in Europe (*31*).

Venom proteins IDDF-1 and IDDF-2 contain a common domain, termed DUF4803, which is highly enriched in genomes of braconid species, including *A. japonica* (*32*, *33*) (fig. S10B). A deep learning-based subcellular localization predictor indicated that most DUF4803 proteins are secreted into the extracellular region of insects and daphnia (fig. S10C), supporting the idea that these proteins are secreted from venom glands and may contribute to parasitism. Our informatic analysis revealed 70 DUF4803 domain genes in the *A. japonica* genome and identified 13 DUF4803 domain proteins among 63 candidates (Data S1). Except for *IDDF* genes, RNAi for the other DUF4803 genes applied to the RNAi screen did not suppress apoptosis in host wing discs. This result implies that DUF4803 proteins have functions other than IDD activity. It is also noteworthy that the DUF4803 gene family is diverse, not only within, but also among braconid species. However, only a few copies of DUF4803 are present in the genome of each non-braconid insect. Understanding how rapid evolution of DUF4803 drives the evolution of parasitic strategies in their hosts is of considerable interest.

## Supporting information

Materials and Methods, Figs. S1 to S11, References

## Acknowledgments

We thank Masako Iida, Akari Kunihisa, Ruriko Fukuda, Anastasiia Staroverova, and Tomoka Nakayama for technical assistance. We also thank Kazuo Takahashi, Masahito Kimura, Tatsushi Igaki, Masayuki Miura, Yuki Ishikawa, Hiroko Sano, Mari Suzuki, Magali Suzanne, Todd Schlenke, Kyoichi Sawamura, Masayoshi Watada, Yukihiko Toquenaga, Masafumi Muratani, the Bloomington Stock Center, the Kyoto *Drosophila* Stock Center in Kyoto Institute of Technology, the Kyorin *Drosophila* species Stock Center, the National Institute of Genetics, the Vienna *Drosophila* RNAi Center, and Developmental Studies Hybridoma Bank for providing reagents; and Shunsuke Furihata, Narumi Shioi, Yoichi Hayakawa, and Yoichiro Tamori, Toru Kuraishi, Masaki Kita, Naonobu Fujita, Hideyuki Goto, Tamaki Yano for and critical comments. We are grateful to all members of the Niwa Laboratory for their insightful discussions and comments on the manuscript. The computational resources were provided by the Data Integration and Analysis Facility of the National Institute for Basic Biology. We would like to thank Steven D. Aird for English language editing.

## Funding

JSPS KAKENHI Grant JP16H06279 (PAGS) JSPS KAKENHI Grant 16K20945

JSPS KAKENHI Grant 18K05670 JSPS KAKENHI Grant 21J10894

JSPS KAKENHI Grant 23K13960 JSPS KAKENHI Grant 24H02297

The Japan Science and Technology Agency (JST)/PRESTO Grant JPMJPR19H6

JST/FOREST Grant JPMJFR2263

Ohsumi Frontier Science Foundation

The program of the Joint Usage/Research Center for Developmental Medicine, Institute of Molecular Embryology and Genetics, Kumamoto University K21-09, K22-10, K23-06, K24-15

## Author contributions

Conceptualization: TKa, YSN, RN Methodology: TKa, AT, TS, NM, SK

Investigation: TKa, YSN, HM, NT, HT, MF, MK, TKu, KS, AN

Visualization: TKa, YSN, RN

Funding acquisition: TKa, YSN, RN

Project administration: YSN, RN

Supervision: YSN, RN

Writing – original draft: TKa, YSN, RN

Writing – review & editing: TKa, YSN, HM, NT, HT, MF, AT, MK, TKu, KS, NM, TS, SK, AN, RN

## Data and materials availability

The number and value of samples in each graph of Figures is listed in Data S3. Raw RNA-seq data shown in Fig. 3H and fig. S6D are available from the DNA Data Bank of the Japan Sequence Read Archive (accession number DRR550214–DRR550219 and DRR550220– DRR550231, respectively). *Asobara japonica* genome and gene annotation data are available from the Zenodo (DOI: 10.5281/zenodo.11084901). The raw data of proteomics is deposited in jPOST (https://jpostdb.org). The accession numbers are PXD052039 for ProteomeXchange and JPST003083 for jPOST.

## Supplementary Materials

Materials and Methods

Figs. S1 to S11

References (*34–45*)

Movies S1.

Data S1 to S5

