## Supplementary material for "Parasitoid wasp venoms degrade *Drosophila* imaginal discs for successful parasitism": Materials and Methods, Figs. S1 to S11, References

**The PDF file includes:**

Materials and Methods

Figs. S1 to S11

References (*34-46*)

**Other Supplementary Materials for this manuscript include the following:**

Movie S1

Data S1 to S5

**Materials and Methods**

Biological materials

For the endoparasitoid wasp, *Asobara japonica*, the parthenogenetic strain "*Tokyo (TK*)" was used32. For the fruit fly, *Drosophila melanogaster*, the *Oregon R (OR*) strain was used as the wild type and host. For *D. elegans*, *HKG-MTK* strain (E-13202, a gift from Yuki Ishikawa, Nagoya University, Japan) was used. A strain of *D. ficusphila* was obtained from the Ehime Fly Stock Center (a gift from Masayoshi Watada). All experimental animals were reared on standard cornmeal-yeast-agar medium at 25 °C under a 12:12 h light/dark cycle. In all experiments, young adult female wasps (aged 1–5 days after eclosion) were used.

The following fly strains were used: *C765-GAL4* (BL36523, a gift from Tatsushi Igaki, Kyoto University, Japan); *WP-GAL4* (BL49828, a gift from Masayuki Miura, The University of Tokyo, Japan) (*34*); *UAS-mCD8::GFP* (BL32186); *Cg-GAL4* (BL7011); *Hml-GAL4>UAS-GFP* (BL6397); *UAS-p35* (BL5072); *reaper*$^{87}$ (BL83150);*UAS-GC3Ai* (a gift from Magali Suzanne, Université de Toulouse, CNRS/UPS, France)(*15*); *UAS-mCherry::Atg8a* (BL37750); *UAS-WT dRubicon-HA* (a gift from Mari Suzuki, Tokyo Metropolitan Institute of Medical Science, Japan) (*21*); *EGFP-Vasa* (a gift from Satoru Kobayashi, University of Tsukuba, Japan) (*35*).

Wasp infection experiment

Wild-type *Drosophila* eggs were laid on grape juice agar plates containing yeast paste for 24 h. Newly hatched first-instar (L1) larvae were transferred to small food vials. At the appropriate developmental stage, fly larvae and adult female wasps were transferred to petri dishes (3–6 cm in diameter) containing a piece of grape agar and cornmeal food. Using a dissection microscope, we observed oviposition behavior of individual wasps and paralyzed host larvae, as described previously (*36*). After female wasps withdrew their ovipositors, paralyzed host larvae were transferred to new, small food vials using forceps. Host larvae recovered within 10 min after infection and grew normally into pupae. After eclosion, numbers of adult wasps, adult flies, pharate adult wasps, pharate adult flies, and dead pupae were counted to calculate parasitism success rates. Uneclosed pupae were considered cases in which parasitism proved lethal to fly larvae.

Microinjection experiment

Whole bodies of 10 female wasps were homogenized with a pestle in 100 μL PBS in a 1.5-mL disposable homogenizer tube (#320103, Nippi). Venom glands and ovaries were dissected from 10 wasps and homogenized separately in these tubes. Samples were centrifuged at 14,000 rpm for 5 min at 4 ºC to collect supernatants. Supernatants were colored blue by adding a food dye, Erioglaucine disodium salt (0.1 g/10 mL Milli-Q) (861146; Sigma). The blue color confirmed that the liquid was injected into the fly larval body. Lysates were injected into wandering L3 fly larvae stuck on glass slides with tape glue under a dissecting microscope. Glass needles were made from a glass capillary filament (GD-1, NARISHIGE) using a puller (PC-10; Narishige). Tips of needles were sharpened using a microgrinder (EG-401; Narishige). Although the precise amount of solution injected could not be controlled because of limitations imposed by our injection apparatus, we estimated that approximately 100–200 nL were injected. Injected fly larvae were transferred to small food vials and were dissected at appropriate times.

Ex vivo co-culture experiment

Venom glands were dissected from 10–20 female wasps in Schneider's *Drosophila* medium (SDM, #21720024; Thermo Fisher Scientific), supplemented with 0.5 mg/mL human insulin

(I9278; Sigma), 10% fetal bovine serum (#10270106) and 5% penicillin-streptomycin (PS) (#26252-94; Nacalai Tesque, Inc.). Fly wing discs were also dissected from wandering L3 larvae in SDM. These samples were co-incubated in glass dishes containing 100 µL SDM on a rotator at 25 ºC for 2 h and then fixed for immunostaining.

Antibody generation

Antibodies against IDDF-1 protein were raised in rabbits and guinea pigs. KLH-conjugated synthetic peptides (NH2-PMRENENDNLPLATAKP-COOH) corresponding to parts of the N-terminal region of IDDF-1 were used for immunization. Antibodies against IDDF-2 protein were raised in rabbits. A KLH-conjugated synthetic peptide (NH2-GELESKQPWMDKFQELLNTT-COOH) corresponding to part of the C-terminal region of IDDF-2 was used for immunization.

Immunohistochemistry

Tissues were dissected in PBS (35501-15; Nacalai Tesque, Inc.) and fixed in 3.7 % formaldehyde in PBS for 30 to 60 min at 25 ºC. Fixed samples were washed 3 times in PBS supplemented with 0.3 % Triton X-100 (PBT), blocked in blocking solution (PBT and 2 % bovine serum albumin [BSA], A9647-1006; Sigma-Aldrich) for 1 h at room temperature, and incubated with primary antibodies in blocking solution at 4°C overnight. Primary antibodies used were as follows: rabbit cleaved Dcp-1 (#9578; Cell Signaling Technology, 1:200), chicken anti-GFP (#ab13970; Abcam, 1:4000), mouse anti-pH3 (#9706; Cell Signaling Technology, 1:400), rabbit anti-Atg8a (ab109364; Abcam, 1:200), guinea pig anti-Shroud (1:1000) (*37*), mouse anti-LacZ (β-galactosidase) (#40-1a; *Drosophila* Hybridoma Bank[DSHB], 1:100), mouse anti-γH2AV (UNC93-5.2.1; DSHB, 1:200), rabbit anti-IDDF-1 (1:1000), guinea pig anti-IDDF-1 (1:200), rabbit and anti-IDDF-2 (1:1000). After washing, fluorophore (Alexa Fluor 488, 546, or 633)-conjugated secondary antibodies (Thermo Fisher Scientific) and phalloidin (A12379, A12380; Thermo Fisher Scientific) were used at a 1:200 dilution. Samples were incubated for 2 h at room temperature in blocking solution. Tissues were then washed again before nuclear staining with 4',6-diamidino-2-phenylindole (DAPI; 62247; Thermo Fisher Scientific, 1:10000) for 15 min. All samples were mounted using FluorSave reagent (#345789; Merck Millipore). Images were obtained with a 10×, 20×, or 40× (water-immersion) objective lens using a Zeiss LSM 700 or 900 confocal microscope. Images were processed using Fiji/ImageJ software (National Institutes of Health, Bethesda, MD).

Lysotracker staining

Tissues were dissected in SDM and cultured in new tubes filled with Lysotracker Red DND-99 (1:250, diluted with SDM; #L7528, Thermo Fisher Scientific) for 30 min. Samples were washed twice with PBS, and fixed for immunostaining.

Dihydroethidium (DHE) staining

Tissues were dissected in SDM and cultured in new tubes filled with 60 μM DHE (#12013, Cayman Chemical Company) for 7 min. Samples were washed twice with SDM, and fixed for immunostaining.

Transmission electron microscopy

Fly tissue samples were fixed with 2% paraformaldehyde (PFA) and 2% glutaraldehyde (GA) in 0.1 M phosphate buffer (pH 7.4) at 4ºC. Wasp tissue samples were fixed with 2% PFA and 2 % GA in 0.1 M cacodylate buffer (pH 7.4) at 4 ºC overnight. After fixation, samples were washed 3 times and postfixed with 2% osmium tetraoxide at 4 ºC for 2 h. Samples were dehydrated in graded ethanol solutions (50, 50, 90, and 100 %) and were infiltrated with propylene oxide (PO). Samples were gradually transferred to a fresh 100% resin (Quetol-812; Nisshin EM Co. Tokyo, Japan) and polymerized at 60 ºC for 48 h. Grids were observed under a transmission electron microscope (JEM-1400Plus; JEOL Ltd., Tokyo, Japan) at an acceleration voltage of 100 kV. Digital images (3296 × 2472 pixels) were captured using a CCD camera (EM-14830RUBY2; JOEL Ltd., Tokyo, Japan). Samples were prepared and observed by Tokai Electron Microscopy, Inc. (Nagoya, Japan).

RNA sequencing (RNA-seq) analysis

Venom glands were collected from 100 wasps, aged 3–5 days, after eclosion. For RNA-seq of RNAi wasp venom gland samples (Fig. 3H), 10 venom glands were isolated from each replicate of each condition. Venom glands and other parts of the bodies were placed separately in 1.5-mL tubes containing TRIzol reagent (Thermo Fisher Scientific, #15596026). All experiments were performed in triplicate. For validation of RNAi (fig. S6C) 10 venom glands were isolated from two-day-old RNAi wasps.

To detect differentially expressed genes, an average of 14 million reads per biological replicate were obtained. We evaluated the quality of raw single-end reads using FASTQC and trimmed 1 base pair from the 3′ end, adaptors, and reads of <20q base pairs in length from raw reads using Trim galore 0.6.4 (Babraham Bioinformatics). Trimmed reads were aligned to the *A. japonica* genome [Kamiyama et al. GCA_026005395.1] using HISAT2 2.1.0. (*38*) Samtools 1.9 (*39*) and StringTie 2.0.6 (*40*) were used to sort, merge, and count reads. We calculated the number of TMM (MM (trimmed mean of M-values)-normalized fragments per kilobase of combined exon length per one million total mapped reads (TMM-normalized FPKM value) with R 3.6.1, Ballgown 2.18.0 (*41*), and edgeR 3.28.0 (*42*), which were subsequently used to estimate gene expression values. All LogFC values and P-values corrected using the Benjamini–Hochberg false discovery rate (FDR) are presented in Data S1 and S2.

Proteomics analysis by mass spectrometry

To prepare venom lysates, 50 venom glands were dissected in PBS and squashed in 50 μL of PBS with a pestle. Samples were centrifuged for 5 min to remove debris. For the venom secretory solution, 50 venom glands were incubated in 20 μL PBS for 2 h, and the supernatant was collected. Triple replicates of both samples were applied to NuPAGE gels (#NP0323BOX, Thermo Fisher Scientific) and stained with CBB (#24590, Thermo Fisher Scientific) for mass spectrometric analysis.

Proteins in the gel were reduced using DTT (Thermo Fisher Scientific), alkylated with iodoacetamide (Thermo Fisher Scientific), and digested with trypsin and lysyl endopeptidase (Promega, Madison, Wisconsin, USA). Resultant peptides were analyzed on an Advance UHPLC system (Michrom Bioresources, Auburn, California, USA) coupled to a Q Exactive mass spectrometer (Thermo Fisher Scientific) with raw data processed using Xcalibur (Thermo Fisher Scientific). Raw data were analyzed against *A. japonica* protein sequences. A decoy database comprising either randomized or reversed sequences in the target database was used for FDR estimation, and the percolator algorithm was used to evaluate false positives. Search results were filtered against a 1% global FDR to ensure a high confidence level.

Comparative genomics

*A. japonica*-specific genes were identified using tBLASTn v2.9.0 analysis (*43*) against the *L. heterotoma* genome (GCF_015476425.1). Amino acid sequences of *A. japonica* proteins were used as queries in the tBLASTn analysis. An expected value (E-value) of < 10-4 was set as the threshold for conservation. *A. japonica* proteins that scored E-value < 10-4 were considered "*A. japonica* specific". Other proteins were considered "conserved." Genomic data for *L. heterotoma* were downloaded from the NCBI Biotechnology Information database (https://www.ncbi.nlm.nih.gov/).

Subcellular localization prediction

SignalP 5.0 (https://services.healthtech.dtu.dk/service.php?SignalP-5.0) (*44*) was used to detect signal peptide sequences in *A. japonica* proteins. Amino acid sequences of *A. japonica* were used as queries. Deeploc2 (https://services.healthtech.dtu.dk/services/DeepLoc-2.0/) was used for the prediction of subcelluler localization of DUF4803 proteins (*45*).

RNAi screen of wasp venom genes required for cell death-inducing activity

Gene knockdown in *A. japonica* was performed as described previously (*9*). We took out *A. japonica* bodies from their host pupal cases using forceps and placed them on a 2% agar plate. Then, dsRNA of the gene of interest was injected into specimens of *A. japonica*. For the first screening, four 1000 ng/µL dsRNAs were mixed (final conc. 250 ng/µL for each) and injected. For the second screening, a single 1000 ng/µL dsRNA was injected. These RNAi wasps were used for infection experiments with five-day-old fly larvae expressing GC3Ai in the wing pouch (*WP>GC3Ai*). Infection of *A. japonica* was confirmed under a dissection microscope. Six hours after infection, larvae were immobilized in a dish filled with Milli-Q water on ice for 10 min. Anterior parts of the infected larvae were imaged to capture GC3Ai fluorescence using a fluorescence dissection microscope (Leica M165 FC). The wing pouch region was selected, and the mean GC3Ai intensity was calculated using Fiji software (https://imagej.net/software/fiji/downloads). DNA primers for gene cloning are listed in Data S4.

Immunoblotting

For venom gland lysates, 10 venom glands were isolated and crushed in 40 μL PBS. Three to four drops of venom fluid leaking from the tip of the wasp ovipositor were collected and added to 10 μL PBS. To detect IDDF proteins, samples were analyzed using an automated western blot analyzer (Abby, Bio-Techne Corporation, Minneapolis, MN, USA). Rabbit anti-IDDF-1 (1:50) and Rabbit anti-IDDF-2 antibodies (1:100) were used as primary antibodies. An anti-rabbit detection module (Bio-Techne Corporation, DM-001) was used for detection.

Production of recombinant IDDF proteins

Expression vectors containing IDDF were generated by Twist Bioscience Inc. In the N-terminal regions of IDDFs, signal peptide sequences were converted to 6xHis sequences. Ribosome binding sequences were added to the upstream of the start codon. Codon usage was optimized for E. coli. DNA fragments of 1,916 bp or 2,117 bp were inserted between the EcoRI and the NotI site of pET21(+) vectors. Recombinant IDDF proteins were produced using the in vitro translation system PUREfrex 2.0 mini (#PF201-0.1, Gene-Frontier, Japan). Following the manufacturer's protocol, bacterial dihydrofolate reductase (DHFR) was also produced as a positive control for the PUREfrex system. Products were detected with CBB staining and western blotting. For the bioassay, these proteins were injected into naive fly L3 larvae. DHFR was injected as a negative control. These larvae were dissected 4 h after injection.

Prediction of IDDF protein structures using AlphaFold 2

We constructed structural models of IDDF-1 and IDDF-2 using AlphaFold2 (v2.3.1) Non-Docker modification setup (https://github.com/kalininalab/alphafold_non_docker) without optional parameters. The "Maximum template release date to consider" parameter in AlphaFold2 was set to the date the prediction was made.

Motif analysis

To identify DUF4803 proteins across arthropods, we downloaded the genome of *Acyrthosiphon pisum* (GCF_005508785.2), *Anopheles gambiae* (GCF_000005575.2), *Aphidius gifuensis* (GCF_014905175.1), *Apis mellifera* (GCF_003254395.2), *Bombyx mori* (GCF_014905235.1), *Cotesia glomerata* (GCF_020080835.1), *Chelonus insularis* (GCF_013357705.1), *Daphnia pulex* (GCF_021134715.1), *Diachasma alloeum* (GCF_001412515.2), *Fopius arisanus* (GCF_000806365.1), *Leptopilina boulardi* (GCF_019393585.1), *Leptopilina heterotoma* (GCF_015476425.1), *Microplitis demolitor* (GCF_026212275.2), *Microplitis mediator* (GCF_029852145.1), *Nasonia vitripennis* (GCF_009193385.2), *Pediculus humanus* (GCF_000006295.1), *Schistocerca gregaria* (GCF_000005575.2), *Tribolium castaneum* (GCF_000002335.3) from the NCBI datasets (https://www.ncbi.nlm.nih.gov/datasets/) and *Drosophila melanogaster* from FlyBase (http://flybase.org). Each amino acid sequence was analyzed using Interproscan (5.42-78.0) to predict domain architecture. The number of DUF4803-containing sequences in each species was determined.

Statistical analysis

All experiments were performed independently at least three times. Sample size depended on the number of independent experiments required for statistical significance and technical feasibility. All statistical analyses were performed in “R (ver. 3.6.1)” and “GraphPad, Prism (ver. 10.2.2)” software.

Particle analysis

Quantification of fluorescence signals was conducted using ImageJ and Fiji software (*46*). For GC3Ai and pH3 signals, a threshold function was used to specify the area of the signals relative to the area of the wing discs. For mCherry::Atg8a signals, a custom-made R code was used to count the number of particles. This R code is provided in Data S5.

### Figure. S1

**A**

| The decelopmental stage of host larvae | Uninfected/ infected | # of animals | larval lethal | pupal lethal | fly eclosed | wasp eclosed | Parasitism success rate (%) |
|---|---|---|---|---|---|---|---|
| 48 hAH | Uninfected | 30 | 0 | 0 | 30 | | |
| (early L3) | infected | 55 | 0 | 1 | 0 | 54 | 98.2 |
| 72 hAH | Uninfected | 67 | 0 | 2 | 65 | | |
| (middle L3) | infected | 65 | 0 | 1 | 0 | 64 | 98.5 |
| 96 hAH | Uninfected | 30 | 0 | 0 | 30 | | |
| (late L3) | infected | 60 | 7 | 31 | 0 | 22 | 36.7 |
| 120 hAH | Uninfected | 20 | 0 | 0 | 20 | | |
| (within 6 h before pupation) | infected | 29 | 0 | 26 | 0 | 3 | 10.3 |

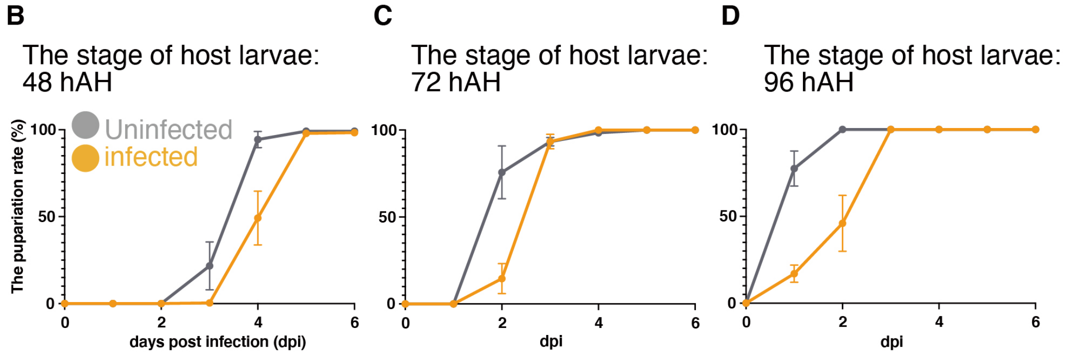


**E**

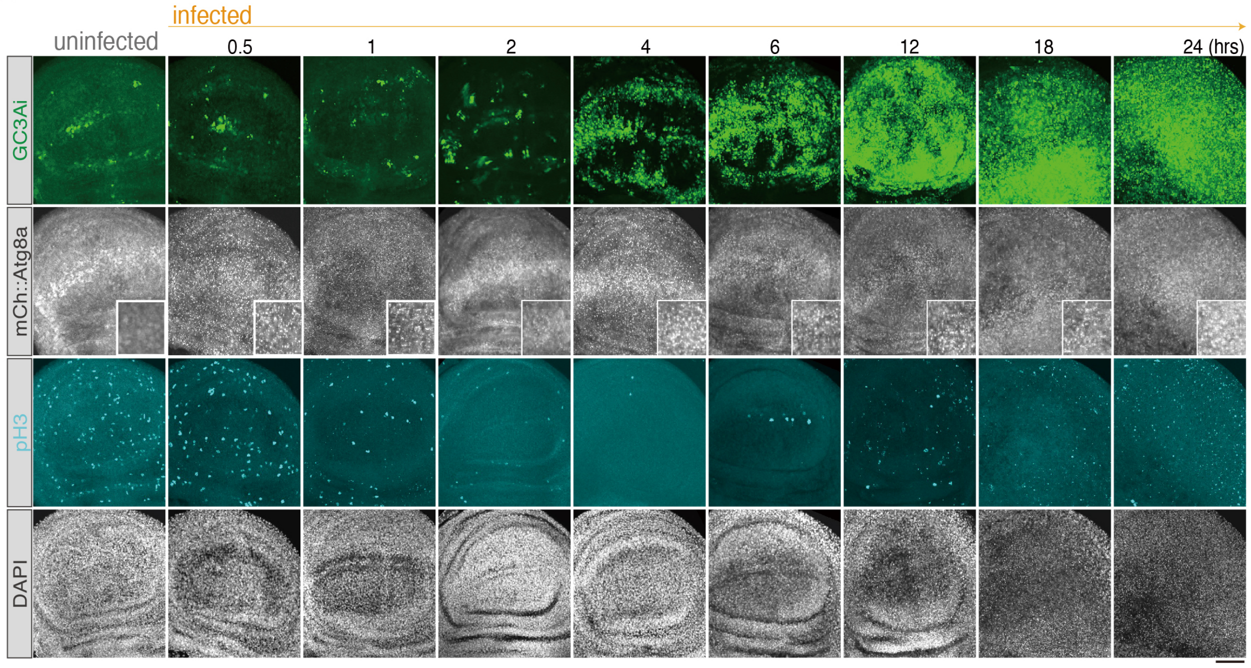


**Fig. S1. The parasitism success rate is related to developmental timing of host larvae in parasitism of the host fly, *D. melanogaster* by *A. japonica*.**

(**A**) The parasitism success rate of *A. japonica* depends on developmental stages of host *D. melanogaster* larvae. When a female wasp deposits an egg in a fly larva during the early to middle third-instar larval (L3) stage (48 to 72 h after hatching, hAH), more than 98% of wasp offspring successfully eclosed from the host pupal case. In contrast, only 37% of wasps eclosed when hosts were in the late-L3 stage. The remaining 63% were dead inside the pupal case. Moreover, 10% of wasps survived when hosts became pupae soon after infection. In any case, flies never eclosed upon infection by *A. japonica*. (**B-D**) Pupariation timing of host fly larvae was delayed one day, compared to that of uninfected larvae during the L3 stage (48, 72, and 96 hAH). In the Y axis, the total number of pupae is set as 100%. (**E**) Time-course analysis of imaginal disc degradation (IDD) after *A. japonica* infection. GC3Ai (green), mCherry::Atg8a (white), pH3 (cyan), DAPI (white) in wing discs of uninfected and infected (0.5, 1, 2, 4, 6, 12, 18, and 24 hpi) larvae. The same images and quantitative analysis are shown in Fig. 1D-G. Insets of mCherry::Atg8a images are magnified views of the dorsal-ventral boundary. pH3-positive cells were visualized using anti-pH3 antibody. Note that autophagy was induced as early as 30 min after infection, which is followed by mitotic arrest around 1 hpi. Apoptosis was prominent at 4 hpi. Scale bar, 50 μm.

### Figure. S2

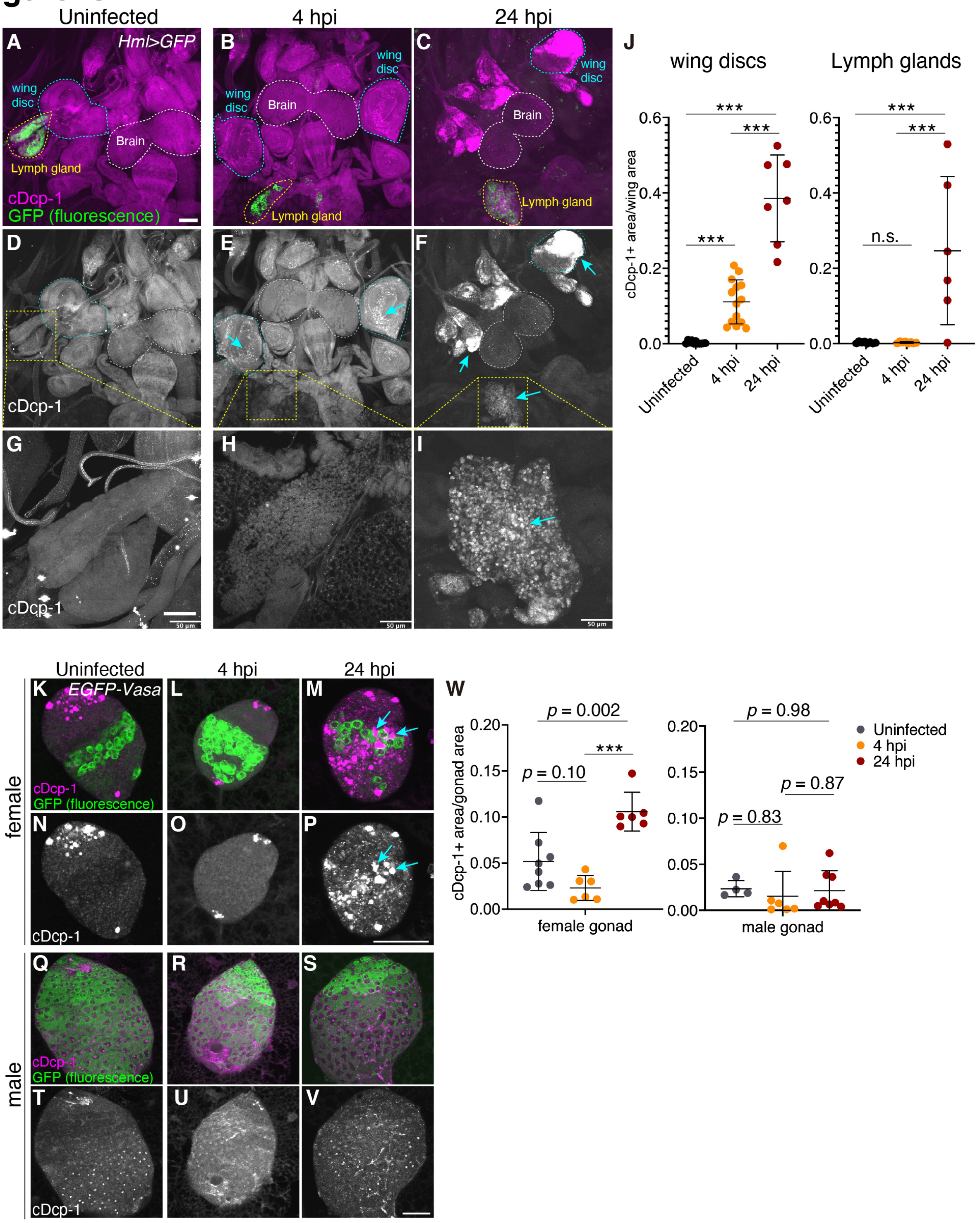


**Fig. S2: Apoptosis is induced in imaginal discs before it occurs in the lymph gland and gonads.**

(**A-I**) GFP (green) and cDcp-1 (magenta in A-C, white in D-I) in the fillet preparation of uninfected and infected (4 and 24 hpi) larvae expressing GFP in the lymph gland (genotype:

*Hml>GFP*). The lymph gland was visualized with GFP. Magnified images of the GFP-positive lymph gland (A-C) are shown in G-I. The cDcp-1-positive area was increased in wing discs at 4 hpi (blue arrows), whereas apoptosis was hardly detected in the lymph gland at 4 hpi (H). At 24 hpi, apoptosis was detected in the lymph gland (I). (**J**) Quantitative analysis of cDcp-1-positive area. (**K-V**) EGFP-Vasa (green) and cDcp-1 (magenta) in the gonads of uninfected and infected (4 and 24 hpi) larvae expressing GFP in germ line cells (genotype: *EGFP-Vasa*). Although apoptosis was detected in some somatic cells even in uninfected larvae, it was hardly detected in GFP-positive germline cells. Scale bar, 50 µm. (**W**) Quantitative analysis of the cDcp-1-positive area in gonads. Statistics: one-way ANOVA followed by Tukey's multiple comparisons test. Scale bar, 50 µm.

### Figure S3

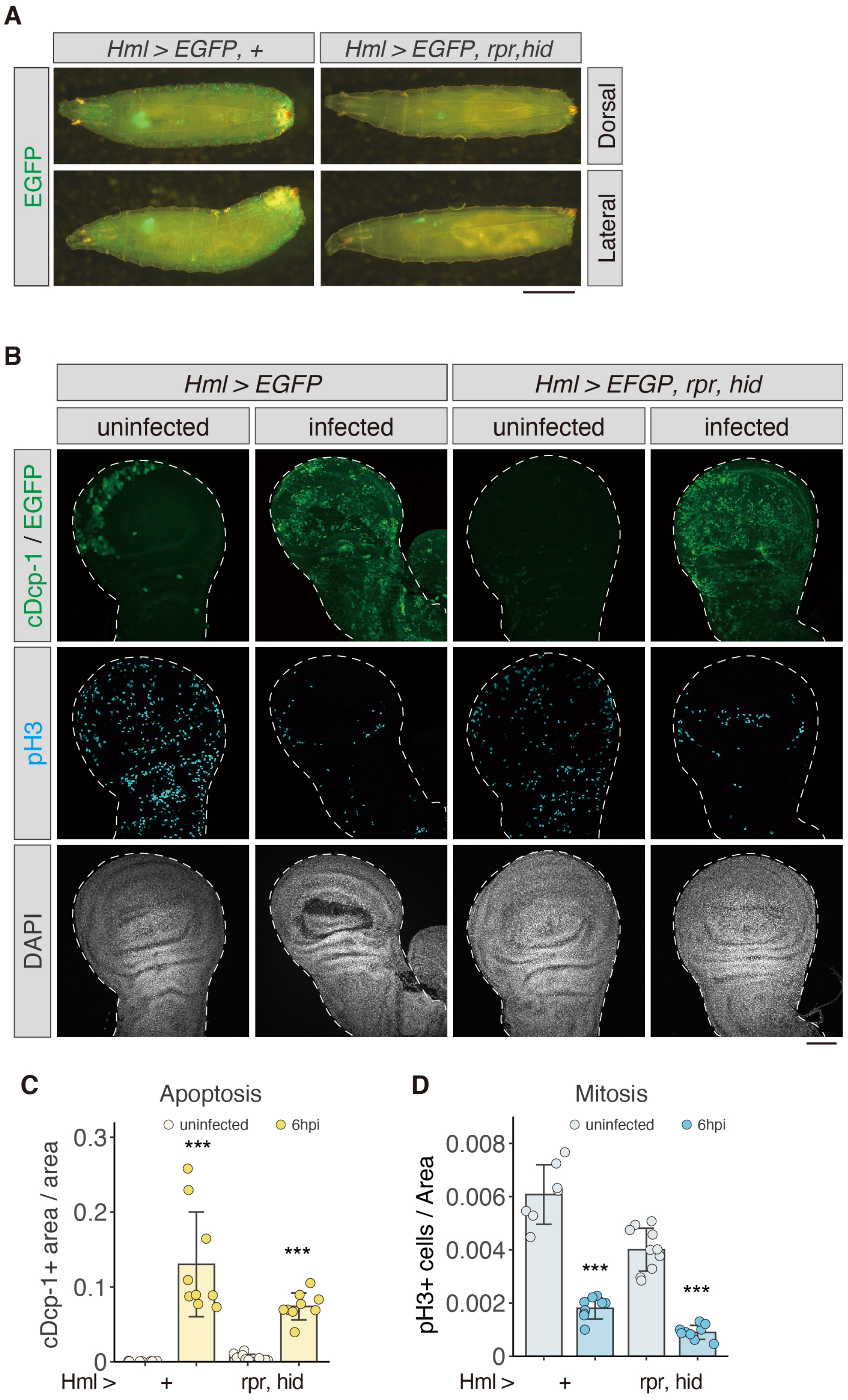


**Fig. S3: Hemocytes elimination did not affect IDD**

(**A**) Dorsal (top) and lateral (bottom) views of L3 larvae expressing *EGFP* in hemocytes. By expressing the apoptotic gene *hid* and *reaper* (*rpr*), most hemocytes were eliminated in larval bodies. Scale bar, 1 mm. (**B**) Wing discs from uninfected and 6 hpi fly larvae with or without hemocyte elimination. cDcp-1 and pH3 were visualized using anti-cDcp1 (green) and anti-pH3 (cyan), respectively. Wing discs were stained with DAPI (white). Scale bar, 50 μm. (**C, D**) Ratio of cDcp-1 positive area and pH3-positive cells to the whole wing disc area ($\mu m^2$) from uninfected and 6 hpi fly larvae with or without hemocyte elimination. Statistics: Two-tailed Welch's t-test between uninfected and 6 hpi samples of each genotype (C, D); *P < 0.05, **P < 0.01, ***P < 0.005; n.s., not significant (P > 0.05).

### Figure S4

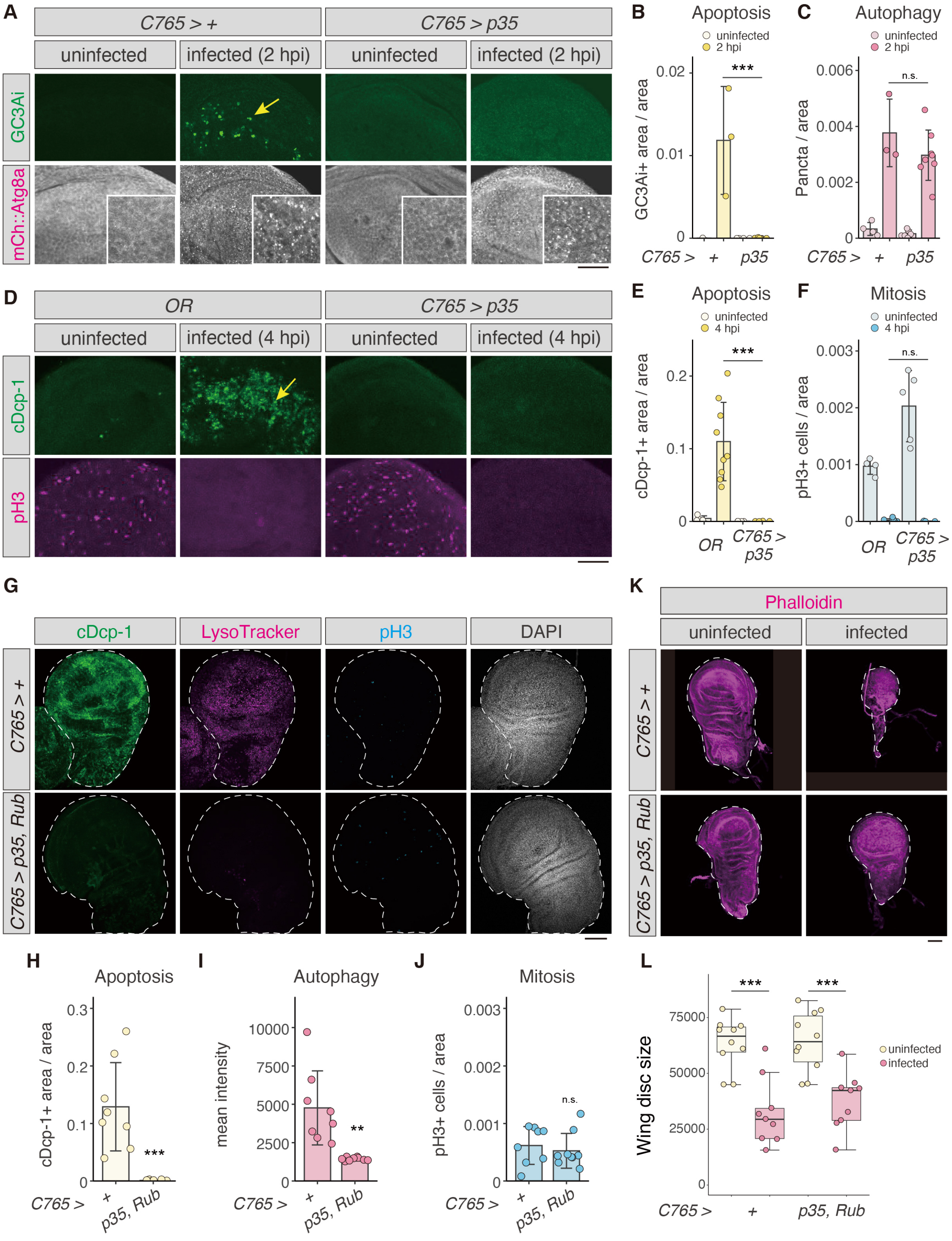


**Fig. S4: Suppression of apoptosis and autophagy did not restore mitosis, resulting in decreased size of wing discs after infection.**

Suppression of apoptosis by expression of *p35* did not inhibit autophagy and mitotic arrest, suggesting that both autophagy and mitotic arrest are independent of apoptosis. In addition, mitotic arrest was still induced in the case of inhibition of apoptosis and autophagy by simultaneous expression of *p35* and *Rub*, indicating that mitotic arrest is independent of both apoptosis and autophagy induction. (**A**) Wing discs from *C765* > + or *C765* > *p35* fly larvae in the uninfected or 2 hpi condition. GC3Ai and mCherry::Atg8a are represented in green and white, respectively. Scale bar, 50 µm. (**B, C**) Quantification of apoptosis (B) and autophagy (C) signals shown in A. (**D**) Wing discs from the wild-type (*OregonR, OR*) or *C765* > *p35* fly larvae in the uninfected or 4 hpi condition. Apoptosis and mitosis are visualized using anti-cDcp1 (green) and anti-pH3 (magenta) antibodies, respectively. Scale bar, 50 µm. (**E, F**) Quantification of apoptosis (E) and mitosis (F) signals shown in D. (**G**) Wing discs from *C765*>+ or *C765* > *p35, Rub* fly larvae at 6 hpi. cDcp-1 and pH3 were visualized using anti-cDcp1 (green) and anti-pH3 (cyan), respectively. Wing discs were stained with LysoTracker (magenta) and DAPI (white). Scale bar, 50 µm. (**H-J**) Ratio of cDcp-1 positive area (H), mean intensity of LysoTracker signals (I), Number of pH3-positive cells per unit area ($\mu m^2$) (J) from uninfected and 6 hpi fly larvae. (**K**) Wing discs of control (*C765* > +) and *p35, Rub*-expressing larvae (*C765* > *p35, Rub*) in the uninfected or infected condition. Larvae of each genotype were infected by *A. japonica* 24 hours after L2/L3 ecdysis and dissected in the prepupal stage. Scale bar, 50 µm. (**L**) Quantitative analysis of wing disc size in each condition. Statistical analysis: Two-tailed Welch's t-test (H and I). Two-tailed Student's t-test (J). One-way analysis ANOVA followed by Tukey's multiple comparison test (b, c, e, f, l). *P < 0.05, **P < 0.01, ***P < 0.005; n.s., not significant (P > 0.05).

#### Figure S5

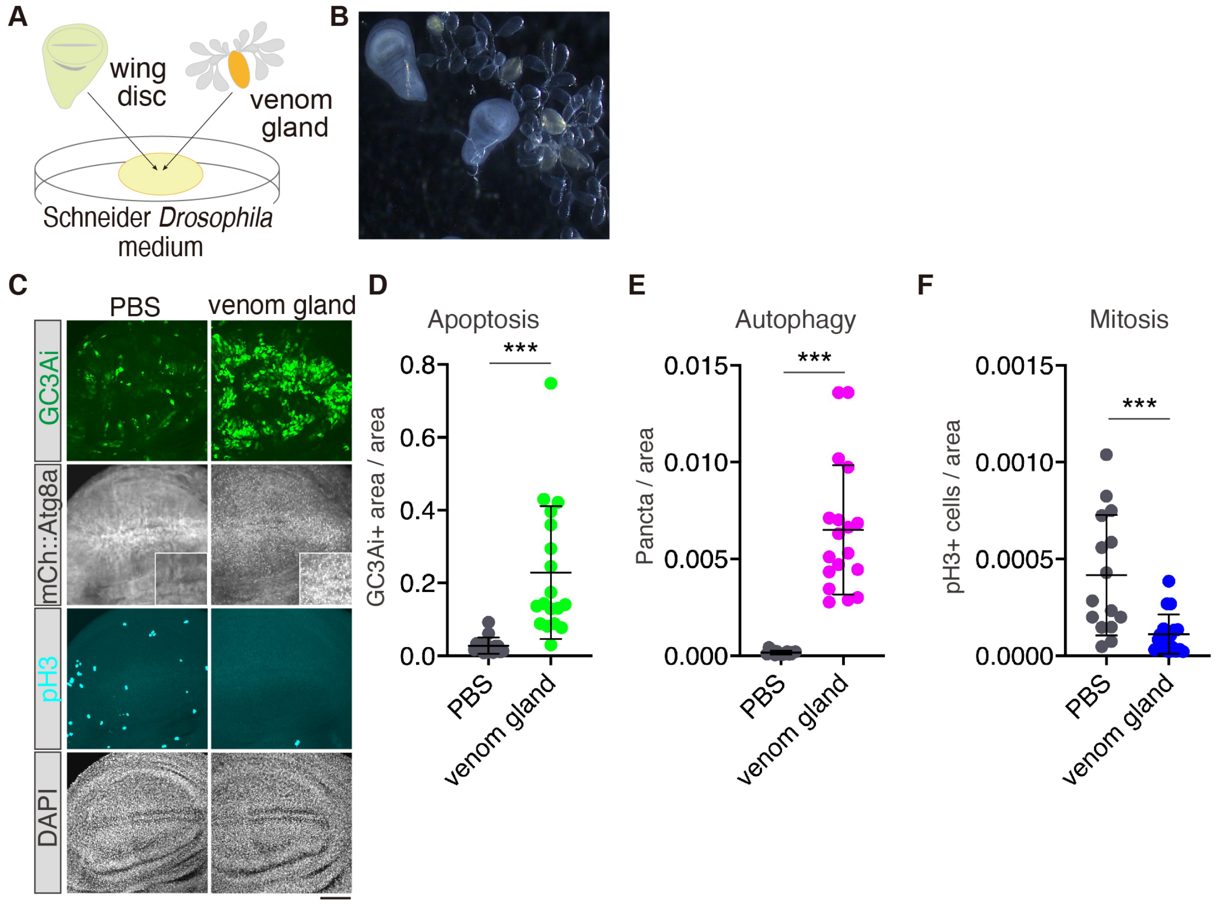


**Fig. S5: IDD-inducing activity is detected by an *ex vivo* co-culture assay with a venom gland of *A. japonica* and wing discs of *D. melanogaster***

(**A**) Scheme of an *ex vivo* co-culture experiment. Two to three pairs of wing discs of *D. melanogaster* and 20 venom glands of *A. japonica* were cultured in 100 µL of Schneider *Drosophila* medium (SDM) for 2 h. (**B**) Two wing discs cultured with venom glands under a light microscopy. (**C**) Cultured wing discs with/without venom glands. GC3Ai (green) and mCherry::Atg8a (white) were detected by their fluorescence. Insets of mCherry::Atg8a images are magnified views of the dorsal-ventral boundary. Mitosis was visualized with anti-pH3 antibody (cyan). Nuclei were stained with DAPI (white). Scale bar, 50 µm. (**D-F**) Quantitative analysis of apoptosis, autophagy, and mitosis in cultured wing discs. GC3Ai signal and mCherry::Atg8a particles were significantly increased, and pH3-positive cells were decreased. Statistical analysis: n=15 (PBS), 18 (venom gland). Two-tailed Mann-Whitney test.

### Figure S6

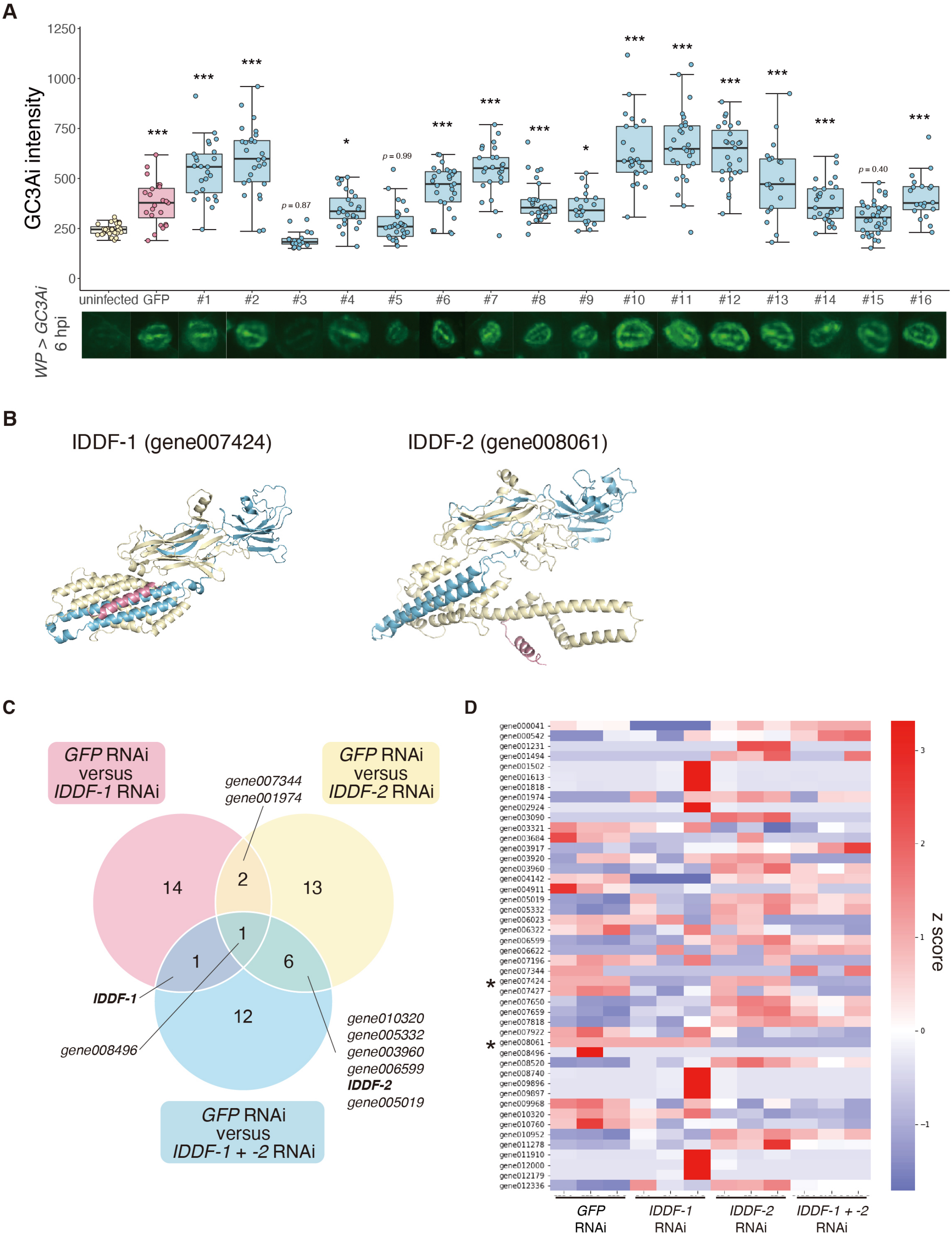


**Fig. S6: RNAi screening of venom genes and predicted 3D-strucures of IDDF proteins**

(**A**) RNAi screening against candidate *A. japonica*-specific genes. Fly larvae expressing *GC3Ai* in the wing pouch region (*WP>GC3Ai*) were used as hosts for RNAi wasps. Fluorescence of GC3Ai increased by infection of control (*GFP*-RNAi) wasps compared to that of uninfected larvae. Box-and-whisker plots representing all biological replicates of GC3Ai intensity. dsRNA mix ID #1~#16 covers 58 candidate target genes. (**B**) Predicted 3D-structure of IDDF-1 (left) and IDDF-2 (right) using AlphaFold2. The signal peptide (SP) and DUF4803 regions are highlighted in red and blue, respectively. (**C**) Venn diagram of differentially expressed genes (DEGs) for *GFP*-RNAi vs. *IDDF-1*-RNAi, *GFP*-RNAi vs. *IDDF-2*-RNAi, and *GFP*-RNAi vs. *IDDF-1* + *-2*-RNAi. FDR < 0.01 is used as a threshold for DEGs. Although *gene008496* was judged as a DEG in all three comparisons, it was probably a batch effect because its expression level was especially high in only one replicate of *GFP*-RNAi sample, for unknown reasons. (**D**) Heatmap of DEGs. Three replicates of Z-scores of DEGs based on FPKMs for each condition were calculated and represented on a color scale (right). Refer to Data S2 for further details.

### Figure S7

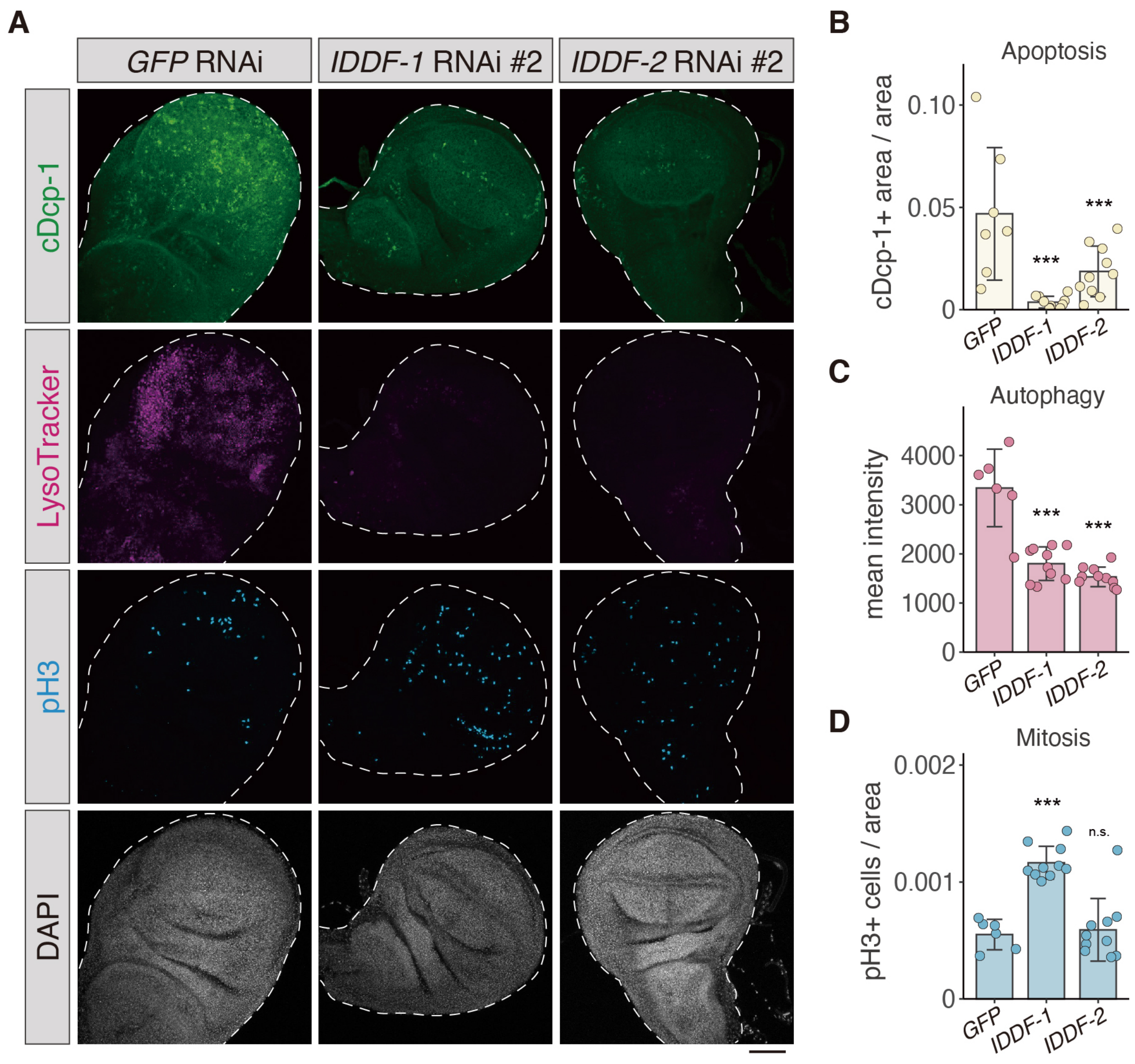


**Fig. S7: The IDD phenotype was suppressed by another target of *IDDF-1* or *IDDF-2* RNAi in *A. japonica***

(**A**) Representative fluorescence images of wing discs of fly larvae infected by *GFP*, *IDDF-1*, or *IDDF-2*-RNAi wasps at 6 hpi. Apoptosis and mitosis were visualized using anti-cDcp1(green) and anti-pH3 (cyan), respectively. Wing discs were stained with LysoTracker (magenta) and DAPI (white). Scale bar, 50 µm. (**B-D**) Ratio of cDcp-1 positive area (B), mean intensity of LysoTracker signal (C) and pH3-positive cells per unit area (µm$^2$) (D) from uninfected and 6 hpi fly larvae were quantified. Consistent with results of Fig. 4C-F, GC3Ai and LysoTracker signals were decreased by *IDDF-1* or *IDDF-2* RNAi whereas mitotic arrest was suppressed only by *IDDF-1* RNAi. Statistical analysis: One-way analysis ANOVA followed by Tukey's multiple comparison test (b-d). *P < 0.05, **P < 0.01, ***P < 0.005; n.s., not significant (P > 0.05)..

#### Figure S8

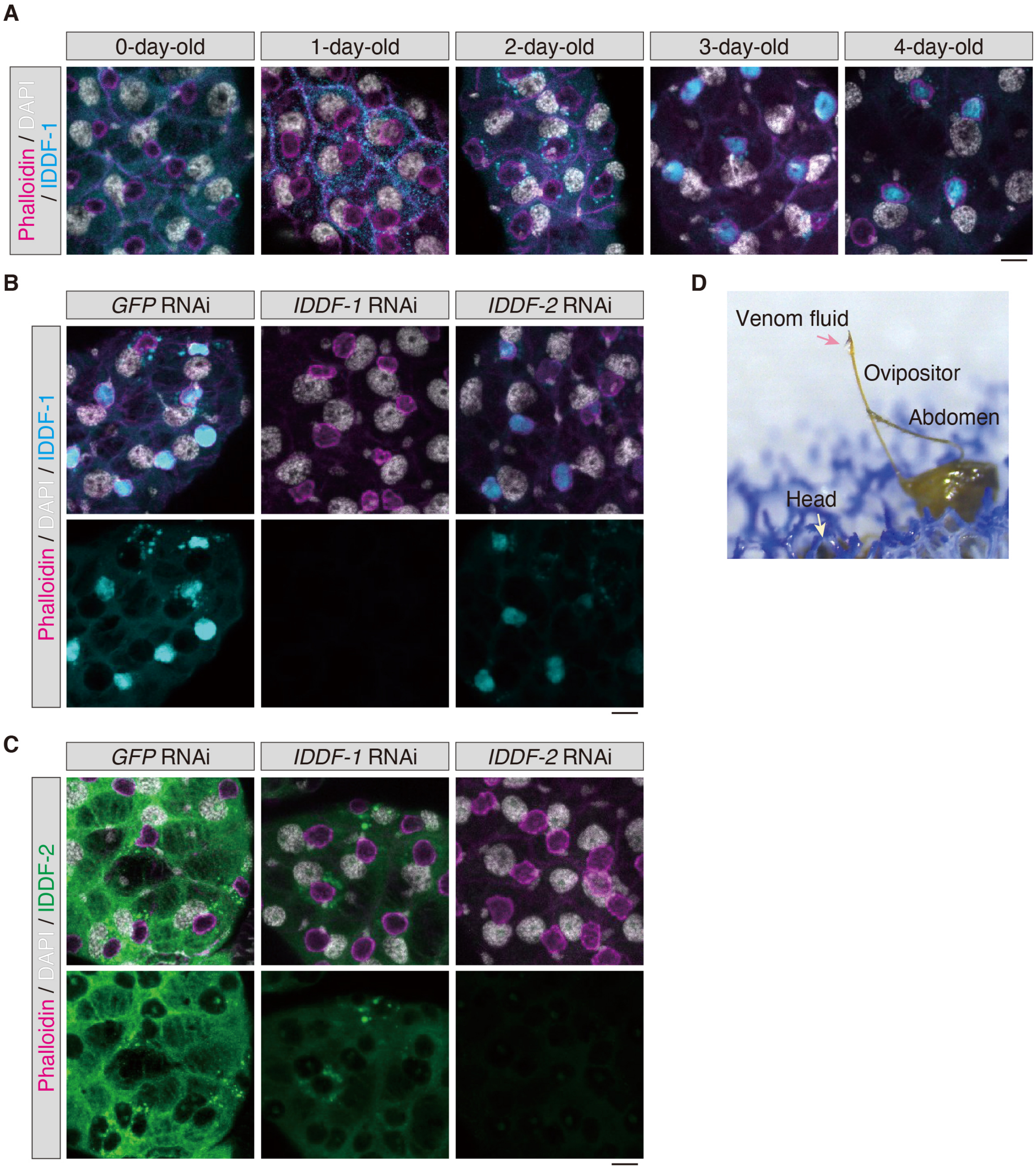


**Fig. S8: Subcellular localization of IDDFs in the venom gland of *A. japonica***

(**A**) Time-course observation of subcellular localization of IDDF-1 (cyan) in the venom gland of *A. japonica*. Samples were stained with phalloidin (magenta) and DAPI (white) to visualize F-actin and DNA, respectively. The size of IDDF-1-containing particles showed an increasing trend days after eclosion. In 3-day-old wasps, the IDDF-1 signal accumulated in the actin-rich ring structures of the venom gland. Scale bar, 10 µm. (**B, C**) Immunostaining of IDDF-1 (B, cyan) or IDDF-2 (C, green) in the venom glands of RNAi wasps. An IDDF-1 or IDDF-2 signal was hardly detected when either of gene was targeted by RNAi. Scale bar, 10 µm. (**D**) Light field view of venom fluid that leaks from the tip of the ovipositor, which was collected for immunoblotting (Fig. 4L). The positions of the wasp head and ovipositor are indicated by white and red arrows, respectively.

### Figure S9

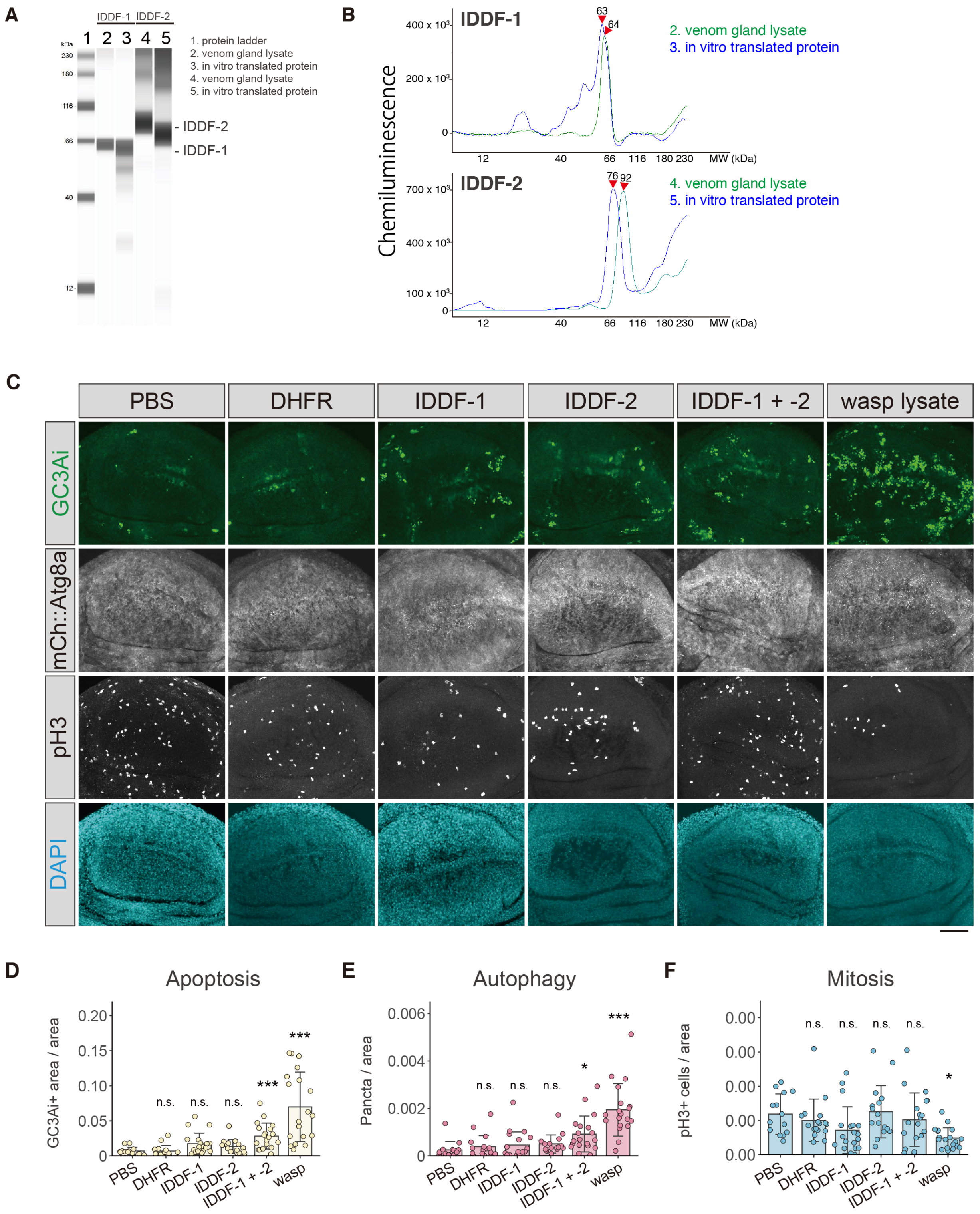


**Fig. S9: In vitro translation of IDDFs and a bioassay with recombinant IDDF proteins.**

(**A, B**) In vitro translation of IDDF proteins detected by western blotting analysis. A protein ladder (1); Venom gland lysate corresponding to a single wasp venom gland (2, 4); IDDF-1

(3); IDDF-2 (5). 1/20 amounts of synthesized proteins were loaded in lanes 3 and 5. The spectrum of peak intensity in immunoblotting is shown in B. Amounts of in-vitro-translated IDDF proteins were comparable to endogenous protein amounts in a single venom gland of *A. japonica*. (**C**) GC3Ai (green), mCherry::Atg8a (white), pH3 (white), and DAPI (cyan) in wing discs from uninfected fly larvae after injection of PBS, dihydrofolate reductase (DHFR, negative control), IDDF-1, IDDF-2, IDDF-1 + -2, wasp lysates. (**D**-**F**) Ratio of GC3Ai-positive area, number of mCherry::Atg8a puncta, number of pH3-positive cells to wing area ($\mu m^2$) of fly larvae in each condition. For all bar graphs, mean ± standard deviation (SD) for all biological replicates is shown. Statistics: Kruskal-Wallis test followed by Dunn's multiple comparisons' test *P < 0.05, **P < 0.01, ***P < 0.005; n.s., not significant (P > 0.05).

### Figure S10

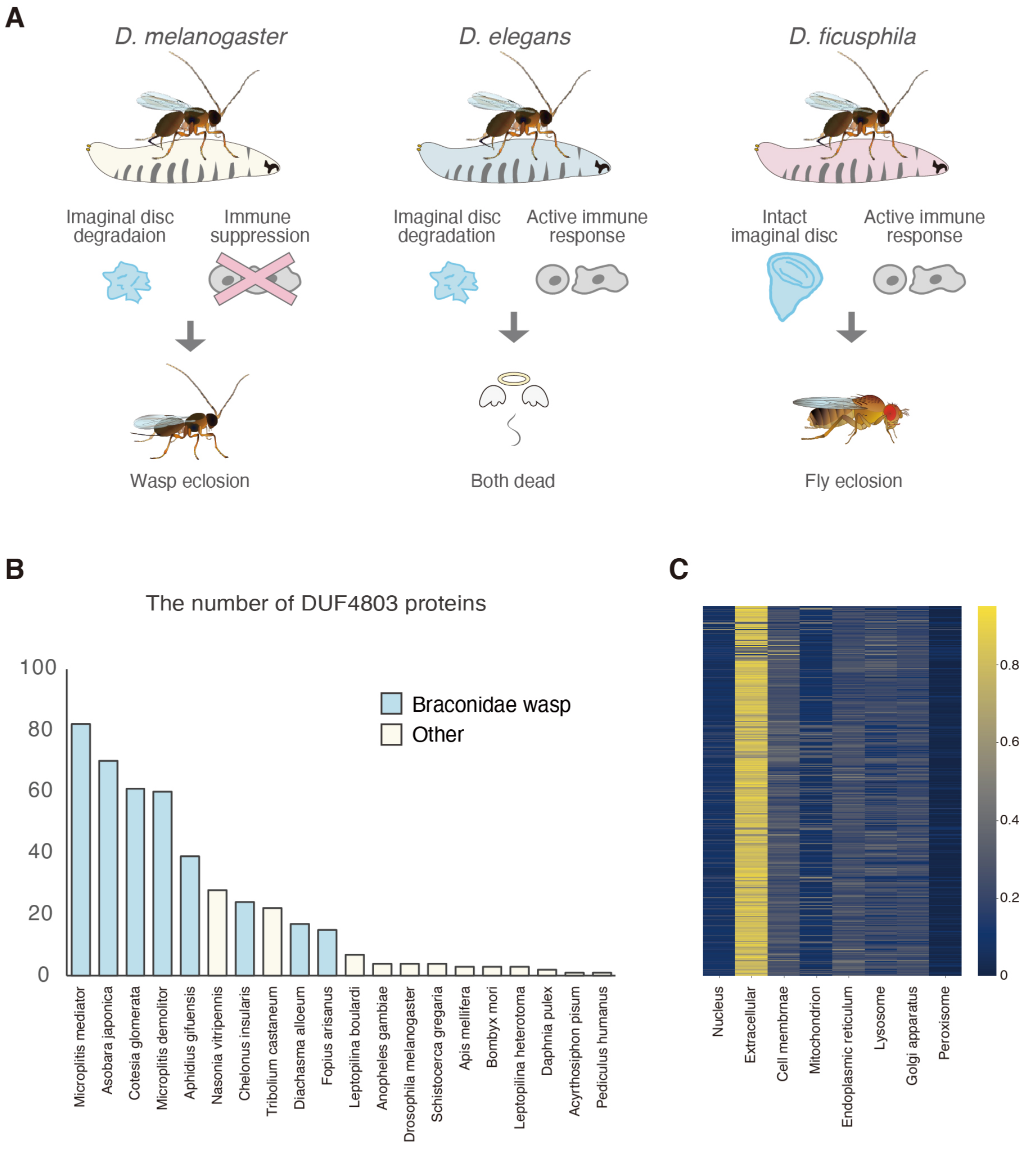


**Fig. S10: The diversity of DUF4803 proteins among arthropods**

(A) Schematics of the response to *A. japonica* infection in *D. melanogaster*, *D. elegans*, and *D. ficusphila*. In contrast to *D. melanogaster*, the immune response is active in *D. elegans* and *D. ficusphila*, preventing successful parasitism by *A. japonica*. (**B**) Numbers of DUF4803 proteins among arthropods. The gene encoding DUF4803-contaning protein is highly duplicated in braconid wasp genomes. (**C**) Probability heatmap of subcellular localization of DUF4803-containing proteins. Probability scores are plotted on a color scale (right side). Most DUF4803-containing proteins are predicted to localize extracellularly.

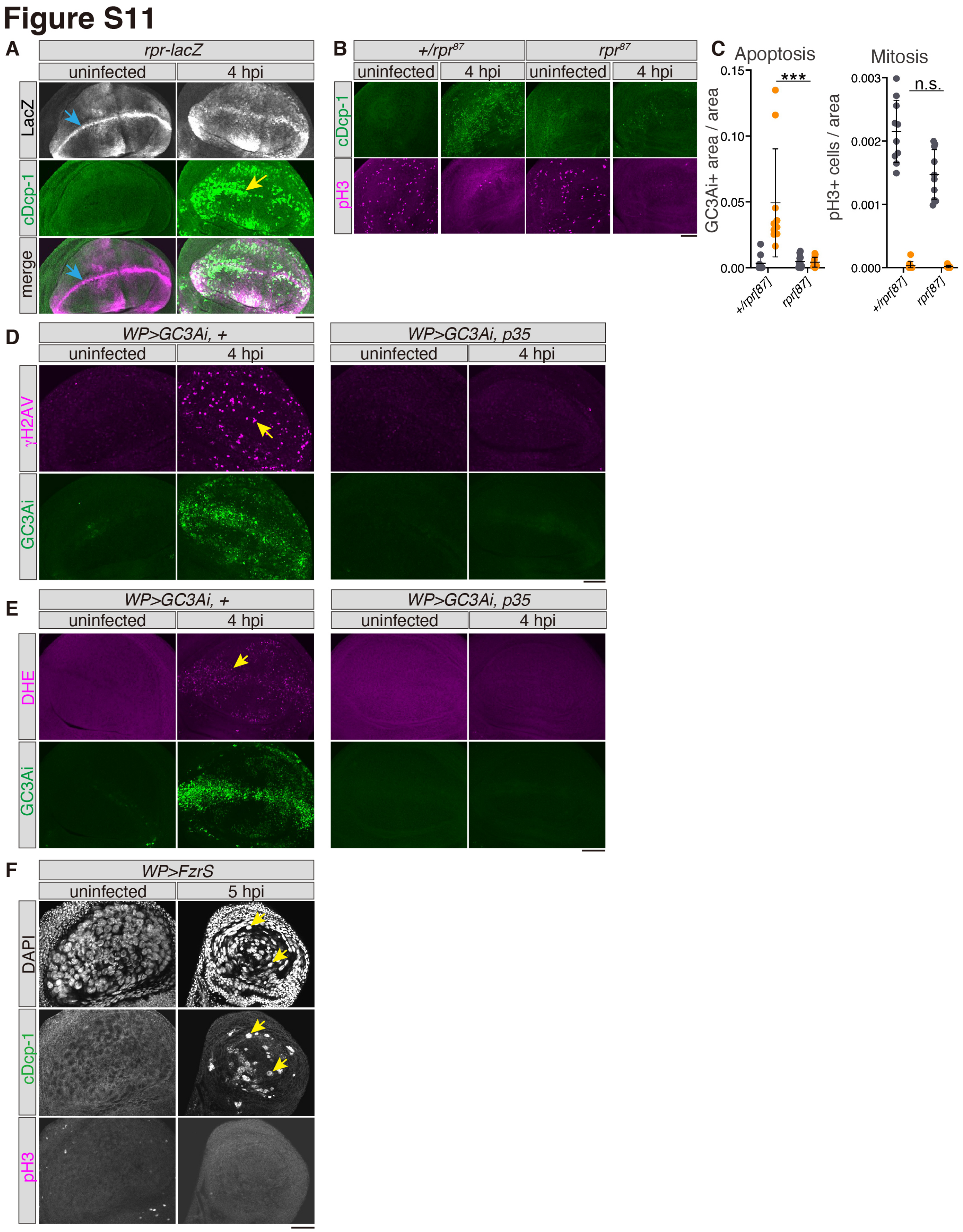


**Fig. S11:** ***Asobara japonica*** **infection-induced apoptosis was mediated by the proapoptotic gene,** ***reaper*****, accompanied by DNA damage and reactive oxygen species.**

(**A**) *rpr-lacZ* was endogenously expressed in the wing pouch region of uninfected fly larvae. Blue arrows indicate strong expression in the dorsal-ventral boundary. Expression of *rpr-lacZ* was upregulated and overlapped with cDcp-1 signals at 4 hpi. Wing discs were stained for LacZ (magenta) and cDcp-1 (green). (**B-C**) *Asobara japonica* infection-induced apoptosis was suppressed in $rpr^{87}$ null mutants. Wing discs were stained for cDcp-1 (green) and pH3 (magenta). Although apoptosis was greatly suppressed, mitosis was not restored in host $rpr^{87}$ mutants. The cDcp-1 signal area (apoptosis, C) and pH-positive cells (mitosis, D) in the wing area were quantified. Statistics: one-way ANOVA followed by Tukey's multiple comparisons test. *P < 0.05, **P < 0.01, ***P < 0.005; n.s., not significant (P > 0.05). (**D**) Wing discs in control *(WP>GC3Ai,* +) and *p35*-expressing larvae (*WP>GC3Ai, p35*). Signal of γH2AV, a marker of DNA damage and GC3Ai increased in wing discs of infected larvae (yellow arrow, 4 hpi), but were hardly detected by *p35* expression. (**E**) Reactive oxygen species (ROS) increased in wing discs of infected larvae at 4 hpi, as indicated by staining with Dihydroethidium (DHE, yellow arrow). DHE signal and GC3Ai signals were hardly detected by *p35*-expression (*WP>GC3Ai, p35*). (**F**) Wing discs in L3 larvae expressing *fizzy-related* (*WP>FzrS*). Overexpression of *FzrS* inhibits mitosis, producing large polyploid cells in the wing pouch region, as indicated by DAPI staining. pH3 signals decreased in uninfected larvae. cDcp-1 signals were detected in these polyploid cells at 5 hpi (yellow arrows). Scale bar, 50 μm.

**Movie S1.** The oviposition behavior of *Asobara japonica*

A female wasp was laying an egg to its host *Drosophila melanogaster* larva on a grape plate agar. The movie was taken by a digital camera (MC120 HD, Leica) attached to a dissection microscope (S8APO, Leica).

**Data S1. Identification of venom gland genes in *Asobara japonica***

Differentially expressed genes (DEGs) represented in Fig. 3H are listed in an Excel file. Results of signalP, proteome analysis and comparative genomics against the *Leptopilina heterotoma* genome are also included. SP represents signal peptide.

**Data S2. Evaluation of the effect of RNAi in *A. japonica*.**

Gene expression profiles of RNAi wasps were analyzed by RNA sequencing. DEGs for *GFP* RNAi vs. *IDDF-1* RNAi, *GFP* RNAi vs. *IDDF-2* RNAi, and *GFP* RNAi vs. *IDDF-1 + -2* RNAi are listed in an Excel file. The results are also shown in fig. S6C, D.

**Data S3. The number and size of samples in each Figure (raw data)**

The number and value of samples in Figs. 1 to 5 and figs. S1 to S11 are listed.

**Data S4. Sequences of primers used in this study**

Primer names and sequences used in this study are listed.

**Data S5. A custom-made R code for counting mCherry::Atg8a particle number**

We utilized the R code to count the number of mCherry::Atg8a particles, representing autophagosomes and autolysosomes.
