## Supplementary material for "Parasitoid wasp venoms degrade *Drosophila* imaginal discs for successful parasitism": Data S5: Data_S5_mCherry-Atg8a_particles_counts_Kamiyama_et_al.R

```
#!/usr/bin/env Rscript
##----------------------------------------------------------------------
## atg8a.R for detecting rashes
##----------------------------------------------------------------------
## all rights reserved to TOQUE, Hitoha Mori, Takumi Kamiyama, Yuko Shimada-Niwa and Ryusuke Niwa
##----------------------------------------------------------------------
## started on 10MAR2023
## last modified on 11MAR2023 for excluding smears
##----------------------------------------------------------------------
## partially depended on the following site
## https://cran.r-project.org/web/packages/imager/vignettes/pixsets.html
##----------------------------------------------------------------------

#install.packages(c("tidyverse","imager"))
#install.packages(c("tidyverse"))
#install.packages("imager")


rm(list=objects())
library(tidyverse)
library(imager)


jpg <- "./data.jpg"

im <- load.image(jpg)
##par(mfrow=c(1,2))

## original
##
plot(im)

readline("hit enter for counting rashes")

## Removing the background noise (gradation)
##
blurSize <- 4
im.bgmod <- (im - isoblur(im,blurSize))
#plot(im.bgmod)

## Binarization
##
im.bin <- grayscale(im.bgmod,drop=T)
thr <- im.bin %>% max * 0.5
im.bin[im.bin >= thr] = 1
im.bin[im.bin < thr] = 0

## Counting and plotting rashes
##
rashes  <- split_connected(im.bin)
cnt <- rashes %>% length
rashes <- rashes %>%
  map(function(rash){
    rash %>% as.cimg
  }) %>% add
rashes %>% highlight(col="skyblue") 
sprintf("no. of rashes = %d",cnt)
```
